# Transposable elements drive reorganisation of 3D chromatin during early embryogenesis

**DOI:** 10.1101/523712

**Authors:** Kai Kruse, Noelia Díaz, Rocio Enriquez-Gasca, Xavier Gaume, Maria-Elena Torres-Padilla, Juan M. Vaquerizas

## Abstract

Transposable elements are abundant genetic components of eukaryotic genomes with important regulatory features affecting transcription, splicing, and recombination, among others. Here we demonstrate that the Murine Endogenous Retroviral Element (MuERV-L/MERVL) family of transposable elements drives the 3D reorganisation of the genome in the early mouse embryo. By generating Hi-C data in 2-cell-like cells, we show that MERLV elements promote the formation of insulating domain boundaries throughout the genome *in vivo* and *in vitro*. The formation of these boundaries is coupled to the upregulation of directional transcription from MERVL, which results in the activation of a subset of the gene expression programme of the 2-cell stage embryo. Domain boundaries in the 2-cell stage embryo are transient and can be remodelled without undergoing cell division. Remarkably, we find extensive inter-strain MERVL variation, suggesting multiple non-overlapping rounds of recent genome invasion and a high regulatory plasticity of genome organisation. Our results demonstrate that MERVL drive chromatin organisation during early embryonic development shedding light into how nuclear organisation emerges during zygotic genome activation in mammals.

## INTRODUCTION

Mammalian genomes are characterised by the presence of a high amount of transposable elements (TEs) with increasingly recognized regulatory features for genome function such as transcription, splicing, and recombination, among others (Feschotte 2008; Bourque et al. 2008; Friedli and Trono 2015; Thompson et al. 2016). In recent years, TEs have also been associated with the three-dimensional organisation of chromatin in the nucleus, such as the inter-chromosomal colocalisation of similar repetitive elements (Cournac et al. 2016), or the occurrence of TEs in domains or at domain boundaries (Dixon et al. 2012; Pope et al. 2014; Darrow et al. 2016; Giorgetti et al. 2016; Winter et al. 2018). This is complemented by evidence for insulator function of specific TE families (Wang et al. 2015; Schmidt et al. 2012). More recently, an association between the expansion of disease-associated tandem repeats and three-dimensional chromatin topology has been reported, highlighting the role of these elements for healthy cellular function (Sun et al. 2018). However, despite their functional relevance, the establishment of a causal relationship between TE genomic location and three-dimensional chromatin organisation has been hindered by the lack of observations in a dynamic system that would allow the examination of changes in chromatin conformation related to TEs.

Interestingly, specific families of TEs display dynamic transcriptional changes during developmental stages around mammalian preimplantation development, allowing the examination of their transcriptional regulatory relevance (Rodriguez-Terrones and Torres-Padilla 2018). This developmental transition involves a remarkably intricate system of tightly orchestrated epigenetic changes that ensure the reprogramming of the terminally differentiated gametes to enable the formation of a totipotent zygote (Burton and Torres-Padilla 2014; Xu and Xie 2018). The characterisation of 2-cell-embryo-like cells (2CLC), derived from mouse embryonic stem cells (mESC), has made it possible to investigate the regulatory mechanisms associated with molecular features of totipotency (Peaston et al. 2004; Macfarlan et al. 2012; Ishiuchi et al. 2015). In particular, recent examinations of 2CLC have hinted at a role for the Murine Endogenous Retroviral Element with a leucine tRNA primer binding site (MuERV-L; also known as MERVL) and the transcription factor Dux in this developmental progression (Hendrickson et al. 2017; De Iaco et al. 2017), highlighting the effectiveness of 2CLC to uncover novel molecular mechanisms related to the establishment of totipotency in vivo. In parallel to epigenetic remodelling of chromatin marks (Wu et al. 2016; Liu et al. 2016; Dahl et al. 2016; Zhang et al. 2016), the three-dimensional organisation of chromatin is rapidly reprogrammed during early mammalian development following zygotic genome activation at the 2-cell embryo stage (Du et al. 2017; Ke et al. 2017). While the molecular mechanisms involved in the establishment of this critical layer of genome regulation in mammals are unknown, work in Drosophila has revealed the establishment of early insulation mediated by the pioneer transcription factor Zelda (Hug et al. 2017), suggesting a role for pioneer factors in the establishment of 3D chromatin during early embryonic development (Hug and Vaquerizas 2018).

In this study, we demonstrate that the chromatin conformation changes that occur during early embryonic development in mouse are driven by the MERVL family of TEs. We first used a new low input Hi-C method to produce high-quality genome-wide chromatin contact maps of mESC and 2CLC. Our data indicate that the chromatin conformation of 2CLC recapitulates features of pre-implantation developmental stages, in agreement with their reported higher level of cellular potency. Unexpectedly, we show that hundreds of MERVL-containing genomic regions in 2CLC undergo 3D chromatin remodelling, specifically gaining domain-insulating properties. We further demonstrate that such structural changes are directly caused by the integration of MERVL elements in the genome, and that structural remodelling occurs in conjunction with the binding of the developmental pioneer transcription factor Dux. Finally, analysis of chromatin contact maps *in vivo* during mouse preimplantation development revealed that the early 2-cell embryo undergoes similar structural changes at MERVL, coinciding with their upregulation. Overall our study provides evidence for a role of repetitive elements in shaping the three-dimensional organisation of the genome and their co-option in regulating key biological functions during early mammalian development.

## RESULTS

### 2CLC display increased three-dimensional structural plasticity genome-wide

To determine the chromatin conformation landscape during the reprogramming of pluripotent mESC towards the totipotent-like 2CLC state, we performed in situ Hi-C in 2CLC. To do so, we first generated 2CLC by depleting the histone chaperone CAF-1 in mESC in vitro, as previously described (Ishiuchi et al. 2015) (Fig. 1a). Briefly, we used an eGFP reporter construct under the control of a MERVL long terminal repeat (LTR), which faithfully identified 2CLC (Macfarlan et al. 2012; Ishiuchi et al. 2015; Rodriguez-Terrones et al. 2018). This allowed us to separate GFP-positive (2CLC) from GFP-negative cells (mESC) using FACS sorting (Fig. 1a and Extended Data Fig. 1a). Immunostaining using an OCT4 antibody and chromocentre inspection confirmed successful separation and purification of 2CLC and mESC (Fig. 1b) (Ishiuchi et al. 2015).

To examine chromatin conformation changes accompanying the mESC to 2CLC transition, we performed *in situ* Hi-C modified for low cell numbers on both types of cells (Fig. 1c-h) with close to a billion sequenced read pairs per sample (Supplementary Table 1) (Díaz et al. 2018). In addition, we compared the obtained mESC and 2CLC Hi-C maps to those from lymphoblastoid cells (Rao et al. 2014), providing a reference for fully differentiated cells. On a whole-chromosome level, chromatin appeared to gain three-dimensional structure progressively, particularly at long contact ranges, as potency levels decrease (Fig. 1c). Indeed, an analysis of A/B compartment strength (Lieberman-Aiden et al. 2009) showed an increase of contacts within the active (A) and inactive (B) compartments, as well as gain of separation between A and B compartments as cellular differentiation progresses (Fig. 1d, I). Further comparisons at the megabase-level showed a similar increase of structure throughout development (Fig. 1e), in agreement with recent observations during mouse neuronal development (Bonev et al. 2017) and the reprogramming of induced pluripo-tent stem cells (Stadhouders et al. 2018). To quantify this effect, we examined the presence of two prominent Hi-C matrix features: topologically associating domains (TADs) -regions of increased self-interaction (Fig. 1f, j) – and loops between genomic regions (Fig. 1g and k). Both TADs and loops showed an increase in their strength in mESC and lymphoblastoid cells compared to 2CLC. Taken together, and in agreement with orthogonal observations of chromatin plasticity using FRAP (Bošković et al. 2014; Ishiuchi et al. 2015), our results suggest that genome organisation increases with developmental.

**Figure 1.**
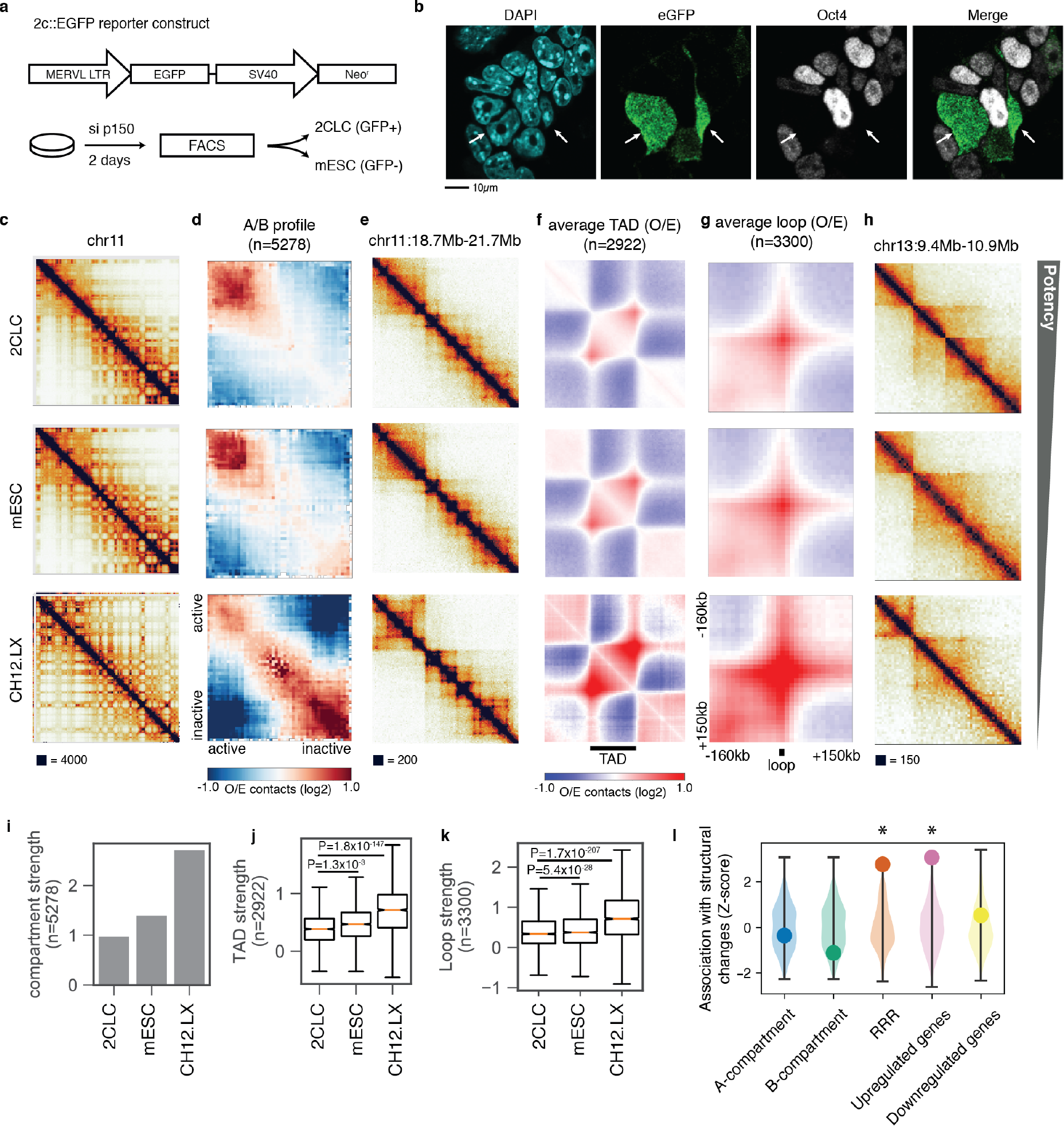
2CLC display genome-wide reduction of chromatin conformation features. (a) 2C::EGFP reporter construct schematic and basic experimental setup for 2CLC induction and sorting. (b) Representative fluorescent microscopy images of GFP−(mESC) and GFP+ (2CLC). White arrows indicate 2CLC. (c-g) Comparison of Hi-C data between pluripotent mESC, 2CLC and lymphoblastoid cells (CH12.LX) (Rao et al. 2014). (c) Chromosome 11 Hi-C maps. (d) A/B profiles showing contact enrichment between active and inactive compartments. (e) Hi-C maps for a 3Mb region on chromosome 11. (f) Observed/expected (O/E) aggregate plot of TADs. (g) O/E aggregate plots of loops. (h) Example of local structural changes observed in 2CLC. (c, e, h) Hi-C contact strength saturation colour indicated as black box underneath the Hi-C maps. (i) Quantification of compartment strength. (j) Quantification of TAD strength. (k) Quantification of loop strength. (j-k) Reported P-values are from Mann-Whitney U test. Boxes span the interquartile range (IQR), i.e. they extend from the first (Q1) to the third quartile (Q3) values of the data, with a line at the median. Whiskers span [Q1 - 1.5 x IQR, Q3 +1.5 x IQR], outliers are omitted (l) Violin plots of the expected association (Z-score) of up- and downregulated genes, reprogramming resistant regions (RRR), as well as A and B compartments with regions undergoing structural changes in 2CLC. Points indicate actual observed association scores; background distributions were obtained from 1000 permutations of the corresponding region locations. Asterisks represent a P-value of 0.05.

To identify specific changes in chromatin organisation during the mESC to 2CLC transition, we performed an analysis of local architectural differences at high spatial resolution (10kb), which revealed ~1500 genomic regions that undergo varying degrees of structural changes in 2CLC (Supplementary Table 2). These regions gain insulating properties, in many cases forming novel topological domain boundaries (Fig. 1h). We find that regions with structural changes are generally dispersed throughout the genome, with no enrichment in either the A (Fig. 1l, Z-score=−0.36, P-value=0.572, permutation test) or B compartments (Fig. 1l, Z-score=−1.13, P-value=0.84). We do, however, find a strong enrichment of structural changes in reprogramming-resistant regions (RRR), genomic regions usually expressed in the 2-cell embryo that resist epigenetic reprogramming following somatic cell nuclear transfer (Fig. 1l and Extended Data Fig. 1b, Z-score=2.76, P-value<0.001) (Matoba et al. 2014). This suggests that the observed changes might be related to the transition from the 2-cell stage to later developmental stages in normal embryos.

Differential gene expression has previously been shown to contribute to the formation of insulating regions during neuronal differentiation (Bonev et al. 2017). An analysis of gene expression changes between mESC and 2CLC revealed that the observed structural changes in 2CLC coincide with the transcriptional upregulation of the underlying regions (Extended Data Fig. 1c). In agreement with this observation, highly upregulated genes in 2CLC are overrepresented at regions with structural changes compared to a random distribution (Fig. 1l, P-value<0.001, Z-score=3.06). However, a significant proportion of upregulated genes did not show signs of structural changes, and a large number of remodelled regions were not associated with differentially expressed genes (Extended Data Fig. 1d). In total, only 101 of the regions showing structural changes contain the promoter of a differentially expressed gene within 30kb. This suggests that genomic features other than single copy genes drive the majority of conformational changes in 2CLC.

### Establishment of *de novo* domain boundaries at MERVL elements

Given the previously described changes in MERVL expression during the mESC to 2CLC transition (Ishiuchi et al. 2015), we sought to determine whether changes in chromatin conformation occurred at the location of MERVL in 2CLC. Visual comparison of mESC and 2CLC Hi-C maps revealed the formation of insulating regions directly at MERVL loci (Fig. 2a). This observation is corroborated by the strong overrepresentation of MERVL elements at regions undergoing structural changes (Z-score > 3, P<0.001, permutation test) (Extended Data Fig. 2a). In addition to the structural changes found at MERVL loci, we observed a strong upregulation of these regions in 2CLC when compared with mESC (Fig. 2a).

To thoroughly investigate the role of MERVL in the chromatin organisation of 2CL cells further, we first performed an exhaustive classification of MERVL TEs into: (i) complete MERVL, consisting of a MERVL-int element flanked by a pair of MT2_MM long terminal repeats (LRT) in a 5’ to 3’ orientation on each side; and, (ii) isolated MT2_MM, termed solo LTRs, which typically arise from a homologous recombination event (Copeland et al. 1983) (Fig. 2b). As TE integrations have previously been shown to vary substantially between mouse strains (Nellåker et al. 2012), we restricted our analysis to those MERVL confirmed to be present in the strain used in this study, 129P2/Ola (Fig. 2b). Aggregate Hi-C maps centred on the TEs clearly showed that the establishment of insulating regions occurs at both complete MERVL and solo LTRs in 129P2/Ola (Fig. 2c), demonstrating that the presence of the LTR is sufficient to drive changes in chromatin organisation. The presence of MERVL at local structural differences in 2CLC prompted us to examine whether MERVL also associated with long-range conformational changes. To examine this, we calculated the average contact enrichment for all pairs of complete MERVL up to 5Mb apart. This analysis revealed a MERVL-specific enrichment of long-range interactions in 2CLC when compared to mESC (Fig. 2d and Extended Data Fig. 2a) (p-value: 3.32×10-13; Wilcoxon signed-rank test). These results demonstrate that MERVL associate with both local and long-range chromatin reorganisation in 2CLC.

**Figure 2.**
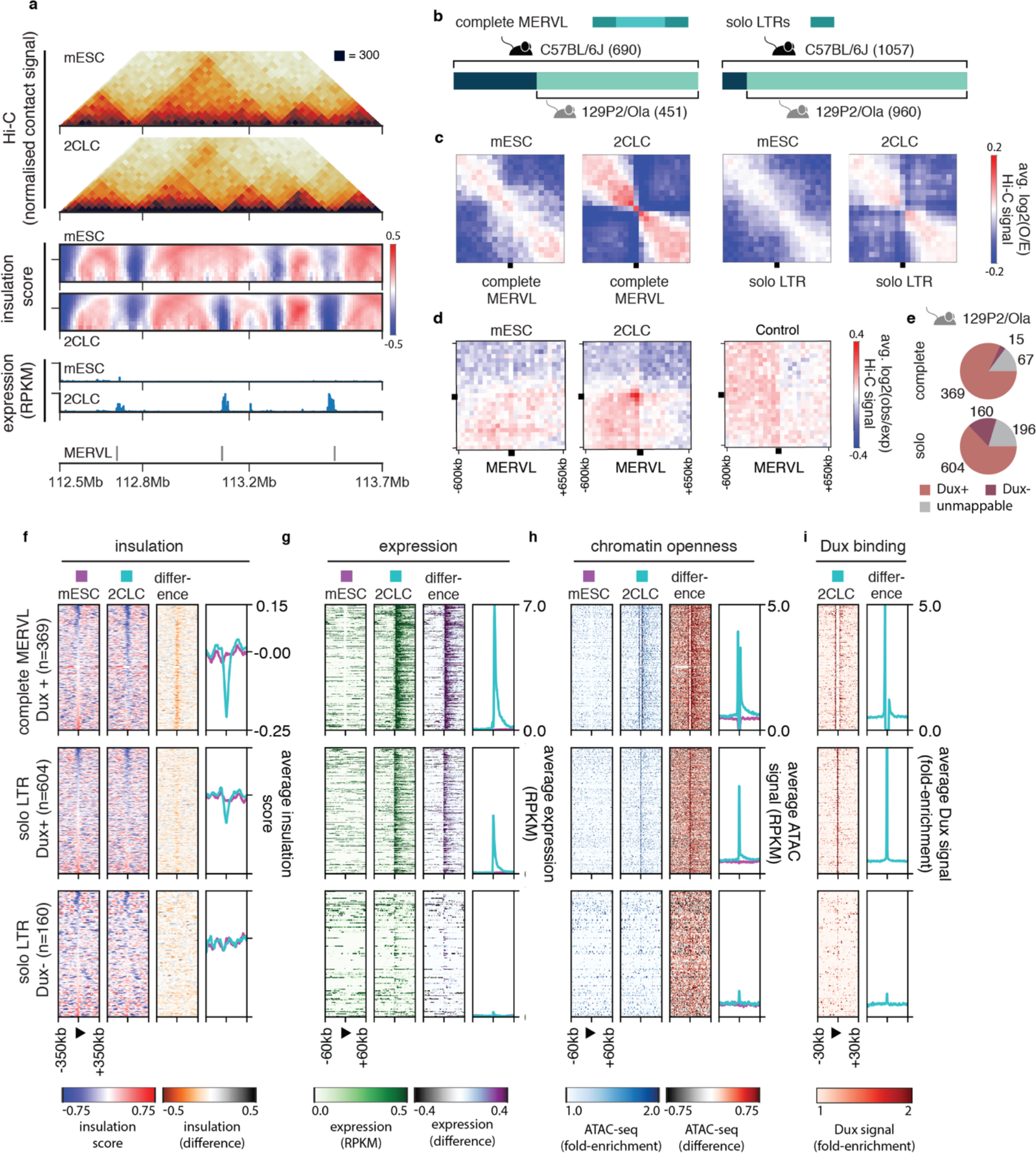
Distinctive 2CLC local and long-range chromatin conformation changes occur at MERVL loci. (a) Representative example region on chromosome 2 highlighting changes in Hi-C, insulation, and expression in 2CLC compared to mESCs at MERVL loci. (b) Schematic representation of complete MERVL and solo LTR, and quantification of MERVL present in strain 129P2/Ola that are annotated in the reference genome of C57BL/6L. (c) Local aggregate Hi-C matrices centred on complete MERVL (left) and solo LTRs (right) in mESC and 2CLC. (d) Long-range aggregate Hi-C observed/expected plots for mESC, 2CLC, and distance-matched control regions in 2CLC for pairs of MERVL regions up to 5Mb apart. (e) Numbers of Dux-bound (Dux+), unbound (Dux-) and unmappable MERVL in the 129P2/Ola genome. (f) Insulation score, (g) expression values (RPKM), (h) ATAC-seq values (RPKM), and (i) Dux binding (fold-enrichment over input) in 129P2/Ola centred on complete MERVL bound by Dux (top), solo LTRs bound by Dux (middle), and solo LTRs not bound by Dux (bottom). Complete MERVL without Dux binding are not shown due to their low numbers (n=15).

We then sought to quantify the extent of structural remodelling occurring at individual TEs using the insulation score (Crane et al. 2015). To do so, we first sub-classified MERVL elements based on their Dux-binding status in 2CLC (positive or negative), since Dux binding is required for MERVL transcriptional activation (Fig. 2e) (De Iaco et al. 2017; Hendrickson et al. 2017). We found that 86% complete, Dux-bound MERVL (284/329, excluding regions with low mappability), and 69% Dux-bound solo LTRs (383/553) displayed gains in insulation in 2CLC (Fig. 2f). Changes in insulation were accompanied by the unidirectional, transcriptional upregulation of the regions downstream of the elements (Fig. 2g). Both expression and insulation changes peak at MERVL and then gradually diminish with increasing distance from the element in a correlated manner (Extended Data Fig. 2b). We next used ATAC-seq data to determine whether changes in chromatin accessibility accompany the structural changes observed at MERVL. This analysis revealed coordinated changes in chromatin accessibility downstream of Dux-bound MERVL elements (Fig. 2h). Thus, we conclude that the changes in chromatin organisation are accompanied by coordinated changes in transcriptional activity and chromatin accessibility at Dux-bound MERVL loci. In contrast to elements bound by Dux, we find that solo LTRs not bound by Dux display no structural changes, downstream expression, or chromatin opening in 2CL cells (Fig. 2f-h, bottom). The number of complete MERVL not bound by Dux (15) was too low to perform similar quantifications, although insulation changes at these loci were not apparent. These results suggest that Dux binding at MERVL is necessary to drive chromatin reorganisation at MERVL loci.

Next, we set to determine whether changes in chromatin organisation were exclusive to MERVL loci, or whether those would appear at other TEs. To address this question, we performed an enrichment analysis at sites of structural changes for all TE families, which revealed a small number of other TEs associated with structural changes in 2CLC (LX4B, BC1_MM, L1MCC) (Extended Data Fig. 2c). An analysis of these elements revealed, however, that the coordinated changes in insulation, expression, and chromatin opening were an exclusive feature of MERVL in 2CLC (Extended Data Fig. 3). In agreement with these results, other families of type I and type II TEs, such as L1 and IAPEZ elements, did not show any signs of *de novo* boundary formation, pervasive transcription or chromatin opening (Extended Data Fig. 4), suggesting that this feature is unique to MERVL.

Finally, we addressed whether Dux binding *per se* was able to induce changes in chromatin organisation. To do so, we examined changes in chromatin structure and accessibility, and transcriptional state in 2CLC for regions bound by Dux not containing an annotated MERVL element. This analysis revealed that while Dux binding in regions of the genome not overlapping with the presence of MERVL elements resulted in an opening of chromatin, Dux binding alone did not result in changes in chromatin conformation or unidirectional pervasive transcription (Extended Data Fig. 4a, b). Overall, these results strongly suggest that binding of Dux alone is not sufficient to result in chromatin architecture re-organisation and that the presence of MERVL elements is necessary for the observed coordinated changes in insulation, expression, and chromatin opening.

### MERVL integration and activation leads to domain boundary formation upon activation in 2CLC

The association between the changes in chromatin architecture and MERVL raises the question as to whether MERVL themselves cause chromatin reorganisation. To test this, we examined the change in chromatin organisation occurring at the genome integration sites of the MERVL LTR-driven eGFP reporter that we used to identify the 2CLC used in Hi-C datasets above, and which recapitulate MERVL endogenous regulation (Ishiuchi et al. 2015). First, we performed virtual 4C analysis using the eGFP sequence as a bait to detect local enrichments in Hi-C interactions across the genome (Díaz et al. 2018). Since Hi-C data have an inherent bias towards detecting interactions between regions located in close regions in the linear DNA sequence, we reasoned that these interactions would identify those regions of the genome where the reporter integrated. This analysis revealed a single interaction peak for the 2C::eGFP reporter on chromosome 12, present in both the mESC and the 2CLC Hi-C data (Fig. 3a, b), suggesting that there is a unique 2C::eGFP reporter integration site in the genome of these cells. An examination of the Hi-C data at the integration site showed a lack of insulation or domain boundary formation in mESC. In contrast, we observed a gain of insulation and the formation of a domain boundary in 2CLC (Fig. 3c,d). These results strongly suggest that the integration of a single MERVL LTR is sufficient to drive changes in three-dimensional conformation at the integration site.

To comprehensively test the ability of MERVL to cause changes in chromatin organisation upon 2CLC reprogramming, we devised an evolutionary analysis strategy that allowed us to test the role of MERVL integrations at many sites in the genome. First, we examined the level of MERVL inter-strain variability using two complementary strategies: (i) an approach based on chimeric reads, using the high-throughput sequencing datasets analysed in this study; and, (ii) a published catalogue of polymorphic TE variants derived from whole-genome sequencing data for 18 mouse strains (Nellåker et al. 2012). We found significant variability of MERVL integrations across all strains (Extended Data Fig. 5a). In particular, both approaches were able to detect a high degree of MERVL integration variability for both complete MERVL and solo LTRs between the reference genome (C57BL/6J) and the strain used to derive the samples used in this study (129P2/Ola) (Fig. 3e, Extended Data Fig. 5a). Incidentally, the absence of C57BL/6J annotated MERVL in the 129P2/Ola genome explains the lack of Dux-binding signal for 239 out of 315 complete MERVL and 109 out of 459 solo LTR classified as not bound by Dux in the 2CLC Dux ChIP-seq datasets, which were produced using cells with a 129P2/Ola background. A close examination of the reads supporting the lack of integrations did not show evidence of target site duplications that usually accompany TE insertions (Extended Data Fig. 5b-f). Together, these results strongly suggest the presence of several independent rounds of MERVL genome invasion during the evolutionary divergence of these strains.

**Figure 3.**
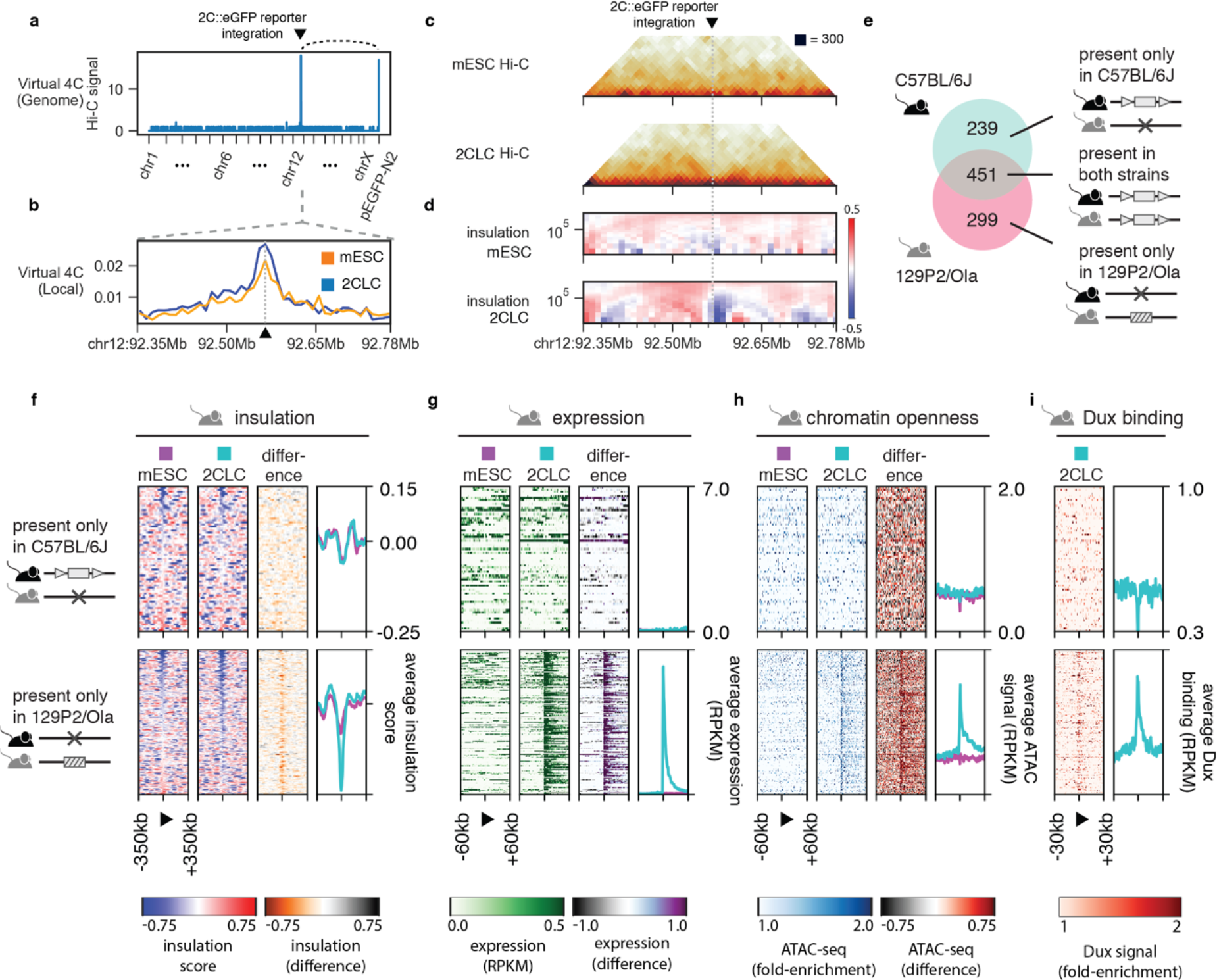
MERVL integration and activation is sufficient to induce structural remodelling in 2CLC. (a) Virtual 4C in 2CLC using the eGFP reporter sequence (pEGFP-N2) as bait. Arc on top highlights interaction peak between the MERVL-driven eGFP reporter and its integration site on chromosome 12. (b) Zoom-in on virtual 4C hit region on chromosome 12, showing reporter integration site in mESC and 2CLC marked by a dashed line and black arrowhead. (c) Hi-C and (d) insulation score plots in reporter integration region for mESC and 2CLC. (e) Venn diagram showing the numbers of complete MERVL specific to C57BL/6J, specific to 129P2/Ola, and shared between the two strains. Datasets in 129P2/Ola showing (f) insulation score, (g) expression values (RPKM), (h) ATAC-seq values (RPKM), and (i) Dux binding (fold-enrichment over input) in 129P2/Ola. (f-i) Heatmaps are centred on complete MERVL only present in C57BL/6J (top), and MERVL integrations only present in 129P2/Ola (bottom).

We examined the ability of MERVL to form domain boundaries genome-wide by analysing changes in chromatin conformation at strain-specific integration sites. To do so, we adapted the chimeric read approach used above to identify missing genomic regions in-between strains, to find novel integrations in the 129P2/Ola genome. Using this method, we detected 299 MERVL integrations in the 129P2/Ola genome that are absent in the C57BL/6J reference genome (Fig. 3e).

We then investigated changes in chromatin insulation, transcriptional activity, chromatin opening, and Dux binding at strain-specific MERVL integrations sites using datasets generated with a 129P2/Ola background (Fig. 3f-i). MERVL integrations not present in 129P2/Ola did not display any changes in chromatin conformation, transcriptional state or chromatin accessibility upon 2CLC induction (Fig. 3f-i, top). These observations confirm that the observed changes in chromatin organisation and transcriptional state upon 2CLC reprogramming cannot be explained by the local endogenous regulatory landscape of these regions before MERVL integration. In contrast, 129P2/Ola-specific MERVL integrations revealed an extensive establishment of insulation, transcriptional activation and increase in chromatin opening at these loci upon 2CLC induction (Fig. 3f-i, bottom). Interestingly, 176 out of 299 of the 129P2/Ola-specific integrations are located at domain boundaries in mESC, which then increase further in insulation in 2CLC (Fig. 3f, bottom). Since strain-specific integrations are likely to be the most recent integrations in the genome, the enrichment of these integrations at domain boundaries, which generally display an active chromatin state (Dixon et al. 2012), suggests that novel MERVL integrations occur preferentially at regions with constitutively active chromatin. We conclude that MERVL element integrations enable the reorganisation of chromatin conformation genome-wide.

### MERVL induce structural changes in early 2-cell stage mouse embryos

Our previous results demonstrate that the Dux-mediated transcriptional activation of MERVL creates novel insulating regions in 2CLC *in vitro*. However, whether a similar reorganisation occurs during early embryonic development is unknown. We thus asked whether MERVL activation at the early 2-cell embryo stage also leads to 3D chromatin re-organisation *in vivo* during this developmental transition (Fig. 4a). To do so, we re-analysed the publicly available Hi-C datasets at 50kb resolution since, due to the scarcity of the data, higher resolution analyses were not possible. Overall, we found very weak signal of broad structural features prior to the 8-cell stage, in agreement with earlier interpretation of these datasets (Du et al. 2017; Ke et al. 2017). However, strikingly, despite the weak chromatin structure present prior to ZGA, we detected the establishment of *de novo* insulation at MERVL loci at the early 2-cell stage (Fig. 4b). Aggregate analysis of the observed/expected matrices at all MERVL locations demonstrated a clear establishment of insulation at these sites, in a remarkable resemblance to those induced in the 2CLC state (Fig. 4c).

A comparison of the insulation score at MERVL elements across the different stages of pre-implantation development (Fig. 4d) revealed that these novel boundaries appear almost exclusively in the early 2-cell stage, and mostly disappear already at the late 2-cell stage. This rapid transition in chromatin conformation is in sharp contrast to previously described changes in chromatin organisation, and demonstrates the existence of fast dynamic chromatin architecture remodelling that does not require passage through mitosis. Interestingly, we found no differences in insulation between MERVL in the *in vivo* datasets regardless of their Dux binding status in 2CLC. Since the early embryo Hi-C experiments were produced using mice with a maternal C57BL/6J background (Du et al. 2017), our results suggest that the entire complement of C75BL/6J MERVL is activated in these embryos. The emergence of TAD boundaries at the early 2-cell stage embryo also coincides with the upregulation of MERVL expression and the presence of pervasive unidirectional downstream transcription in those regions (Fig. 4e). Importantly, an examination of chromatin conformation dynamics for 129P2/Ola-specific MERVL integrations revealed no changes at these locations (Extended Data Fig. 6a). These results strongly suggest that the changes in chromatin organisation *in vivo* are a direct consequence of the MERVL integration rather than the genomic landscape before MERVL integration.

Finally, recent reports have shown that a subset of LINE1 types are also upregulated in mouse early embryonic development (Ancelin et al. 2016; Fadloun et al. 2013; Peaston et al. 2004) and regulate chromatin accessibility (Jachowicz et al. 2017; Percharde et al. 2018). We were therefore interested in determining whether changes in three-dimensional conformation also occur at other TE loci or whether this is a MERVL-specific feature *in vivo*. A heatmap representation of the changes in chromatin conformation and accessibility at these sites showed that the vast majority of elements of each LINE-1 type (L1Md_T, L1Md_Gf, L1Md_A, L1Md_F, L1Md_F3) display no change in these properties (Extended Data Fig. 6b). Therefore, overall, our results demonstrate the presence of MERVL-driven, fast dynamic changes in chromatin organisation at the early 2-cell embryo stage.

**Figure 4.**
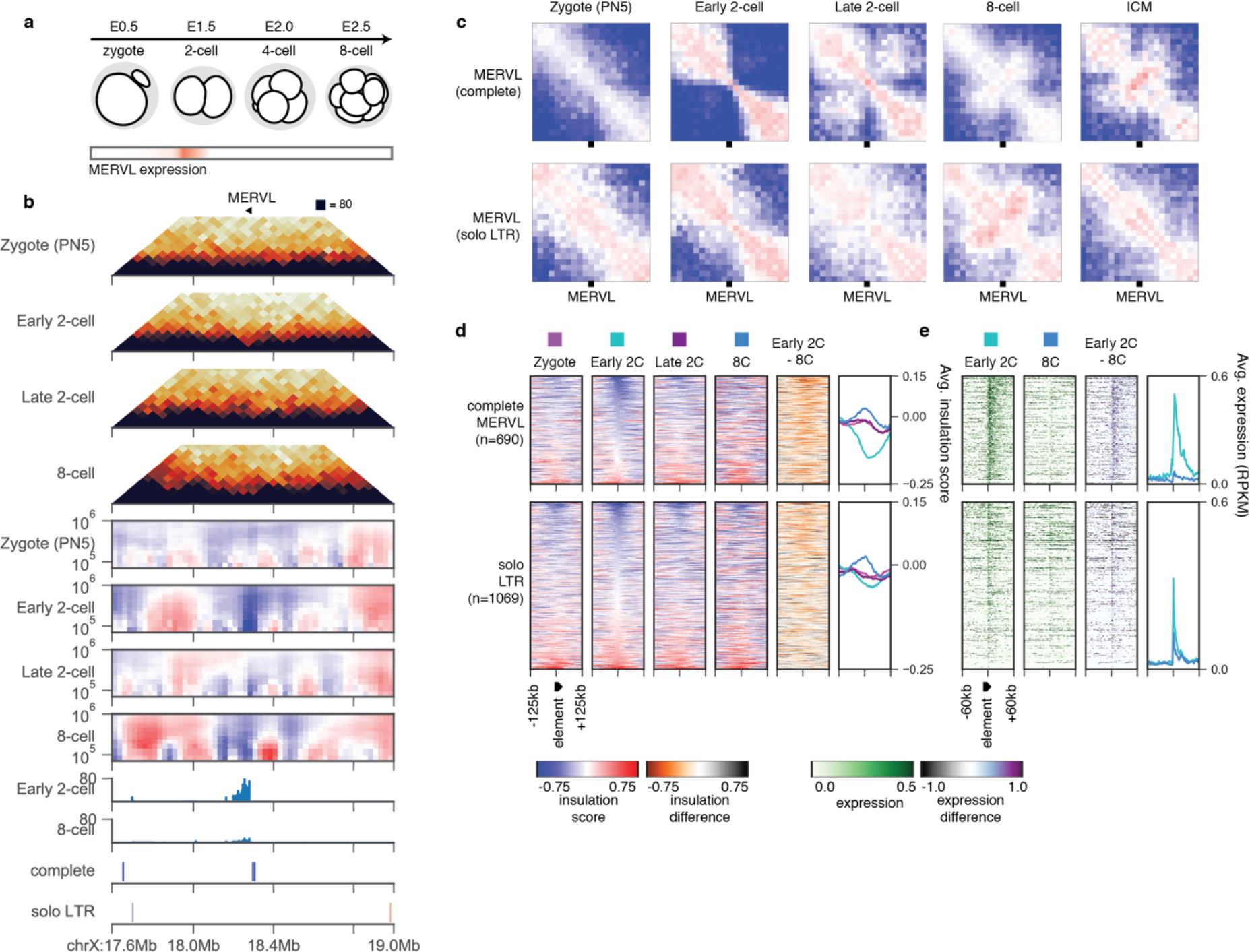
Dynamic establishment of insulation at MERVL in early 2-cell embryos. (a) Schematic overview of mouse early embryonic development with MERVL expression in early 2-cell stage highlighted. (b) TAD boundary formation at MERVL in vivo highlighted by insulation score and expression. (c) Aggregate observed/expected Hi-C plots at complete MERVL and solo LTRs throughout early embryonic development. (d-e) Heatmaps of (d) insulation score and (e) expression for complete MERVL (top) and solo LTRs (bottom) throughout early embryonic development.

## DISCUSSION

Our work has uncovered a novel role for the MERVL family of TEs in driving changes in three-dimensional chromatin organisation. By generating chromatin conformation maps for 2CLC, where MERVL are activated, and comparing these with maps for mESC, we demonstrate that despite the overall high level of similarity between the two maps, striking differences in chromatin architecture appear at hundreds of loci in the genome coinciding with the location of MERVL, characterised by a gain in insulation between neighbouring domains. These results offered invaluable guidance to assist in the examination of the chromatin conformation changes in the early embryo, where the overall level of three-dimensional chromatin organisation is very weak. Importantly, we show that a similar local structural rearrangement occurs at MERVL loci at the early 2-cell stage of early embryonic development. Given the very limited amount of three-dimensional chromatin conformation at this developmental time point, further work is necessary to determine the interplay between the structural changes reported here and the establishment of the complete feature-rich structure found at later developmental stages.

We find that strain-specific MERVL integrations, which are likely to have occurred more recently in evolutionary history, seem to occur preferentially at constitutive TAD boundaries. This is similar to observations made for other TE families, and is generally attributed to a preference to insert into open chromatin regions (Dixon et al. 2012; Pope et al. 2014; Darrow et al. 2016; Giorgetti et al. 2016; Winter et al. 2018; de Jong et al. 2014). On the other hand, MERVL shared between strains, which are likely to have occurred in a common ancestor and thus further back in evolutionary history, are not enriched at TAD boundaries, but can rather be found in a wide range of 3D contexts. This suggests a scenario where regions of ancient MERVL integration were originally constitutively active, and have since evolved into regions where expression, insulation, and chromatin opening are under MERVL control. This would then enable the targeted activation of development-specific transcriptional networks (Macfarlan et al. 2012; Eckersley-Maslin et al. 2016).

In agreement with previous observations during ZGA in *Drosophila* (Hug et al. 2017) and echoing recent observations of TAD boundary formation during cortical neuron development (Bonev et al. 2017), the establishment of TAD boundaries coincides with the recruitment of RNA Pol II and active transcription at those loci. Interestingly, although the two are closely intertwined, the establishment of local insulation is not a general feature of transcriptional upregulation, since many upregulated genes do not undergo changes in chromatin conformation. Since the emergence of domain boundaries is independent of transcription *per se* (Hug et al. 2017; Bonev et al. 2017; Du et al. 2017; Ke et al. 2017), this suggests that the process by which the transcriptional machinery is recruited to these regions, and the associated transcription factors, such as Dux, might play a specific role in the establishment of local insulation.

Taken together, our results demonstrate a role for repetitive elements in dynamically shaping the three-dimensional conformation of chromatin in the nucleus during early embryonic development in mammals. Given the rapid transition between a mostly relaxed architecture during early developmental stages and the establishment of proper chromatin conformation at ZGA, further studies examining the kinetics of this process and its relation to other molecular mechanisms, such as the formation of lamina-associated domains, will shed light into how chromatin is organised during normal mammalian development.

## AUTHOR CONTRIBUTIONS

Conceptualisation: K.K., J.M.V.; Methodology: K.K., N.D., J.M.V.; Software: K.K, R.E.G.; Formal analysis: K.K. and R.E.G.; Investigation: K.K., N.D., R.E.G., J.M.V.; Resources: R.E.G., X.G., M.-E.T.-P.; Writing-Original Draft: K.K, J.M.V.; Writing – Review and Editing: K.K., M.-E.T.-P., J.M.V.; Supervision: J.M.V.; Funding acquisition: J.M.V.;

## ACKNOWLEDGEMENTS

This research was funded by the Max Planck Society. We thank Ivan Bedzhov for critical reading of the manuscript. R.E.-G. was a member of the International Max Planck Research School – Molecular Biomedicine. J.M.V. is a member of the DFG Cells-in-Motion Cluster of Excellence (EXC 1003 – CiM), University of Muenster, Germany. M.E.T.-P. acknowledges funding from ERC-Stg NuclearPotency (280840) and the Helmholtz Association.

## SUPPLEMENTARY INFORMATION

### Methods

#### Cell culture and transfection

E14 mouse ES cells were cultured and as previously described (Ishiuchi et al. 2015). For information on plasmid construction and generation of the stable cell lines please refer to (Ishiuchi et al., 2015). After the p150 knockdown, cells were pelleted (1,000 rpm at 4°C) and resuspended on PBS to be FACS-sorted by GFP expression. FACS Calibur (BD Biosciences) was used to quantify the GFP-positive/GFP-negative cell populations. FACSAria II (BD Biosciences) was used to collect GFP positive and negative cells for the Hi-C experiments.

#### In situ Hi-C library generation for low numbers of cells

We performed in situ Hi-C as described before (Díaz et al. 2018) adapted for cultured cells using a starting material of 100k cells. Data for two biological replicates were generated per sampling point. Briefly, after FACS sorting, cells were cross-linked on a 1% final concentration (v/v) of 37% Formaldehyde and incubated with gentle rotation (20 rpm) at room temperature for 10 min. The quenching of the reaction took place at room temperature with gentle rotation (20 rpm) by adding 2.5M Glycine solution to a final concentration of 0.2M for 5 min. Cells were pelleted twice (300 g, 4°C for 5 min) and resuspended in 1 ml of cold 1X PBS. Cells were pelleted and gently resuspended in 250 μL of ice-cold in situ Hi-C lysis buffer (10 mM Tris-Cl pH 8.0, 10 mM NaCl, 0.2% IGEPAL CA-630, cOmplete Ultra protease inhibitors) and incubated 15 min on ice. Cells were then spun down (1000 g, 4°C for 5 min) and pellet was resuspended in 125 μL of ice-cold in situ Hi-C lysis buffer. Lysed cells were always flash-frozen in liquid Nitrogen and stored at −80°C. Aliquots from the freezer were directly placed on ice, then spun (300 g for 5 min at 4°C) and resuspended in 250 μL in situ lysis buffer. After another spun (13,000 g for 5 min at 4°C), cells were gently resuspended in 250 μL ice-cold 10x NEB2 buffer. Nuclei were then permeabilised by resuspending them in 50 μL of 0.4% SDS at 65°C for 10 min without agitation. SDS was quenched by adding 25 μL of 10% Triton X-100 and 145 μL of nuclease-free water, respectively, at 37°C for 45 min with rotation (650 rpm). Chromatin digestion was done by adding 100U of MboI in 20 μL of 10x NEB2.1 buffer, respectively (New England Biolabs). All the digestions were performed at 37°C with gentle rotation for a period of 90 min by adding the restriction enzyme in two instalments. MboI heat-inactivation was always done post-digestion for 20 min at 62°C. Restriction enzyme generated overhangs were filled-in by adding a mix of 0.4 mM biotin-14-dATP (18.75 μL; Life Technologies), 10mM dCTP, 10mM dGTP and 10mM dTTP (0.75 μL of each dinucleotide), and 5U/μL DNA polymerase I Klenow (New England Biolabs), followed by a 90 min incubation at 37°C with gentle rotation. A master mix containing nuclease-free water (657μL), 10X T4 DNA ligase buffer (120 μL), 10% Triton X-100 (100 μL), 20mg/ml BSA (12 μL) and 5 Weiss U/μL T4 DNA ligase (5μL in two installments; Thermo Fisher Scientific) was added to the samples to ligate the DNA fragments. Samples were mixed by inversion and incubated for 4h at 20°C with gentle rotation. The nuclei were spun down (2,500 g for 5 min at room temperature) and resuspended in in 500 μL extraction buffer. Proteins were digested by adding 20 μL of 20mg/ml proteinase K (Applichem) to the mix and incubating the solution for 30 min at 55°C with rotation (1000 rpm). Afterwards, 130 μL of 5M sodium chloride was added and was then incubated overnight at 65°C with shaking (1000rpm). DNA was precipitated following a Phenol-Chloroform-Isoamyl alcohol (25:24:1) protocol. GlycoBlue (Life Technologies) was used to dye the pellets. Pellets were washed with cold 70% ethanol and all ethanol traces were removed by air drying the pellet for no longer than 5 min. Pellets were dissolved in 30 μL of Tris pH 8.0 (Applichem) and incubated for 5 min at 37°C without rotation. RNA was removed by adding 1 μL RNAse to each sample and incubating the mix 15 min at 37°C. Biotin from unligated fragments was removed by adding 10 μL of 10X NEB2 buffer (New England Biolabs), 1mM of a dNTPs mix (2.5 μL), 20mg/ml BSA (0.5 μL), nuclease-free water (up to 100 μL) and 3U/μL T4 DNA polymerase (5 μL; New England Biolabs). Samples were mixed by gentle pipetting up and down and incubated at 20°C for 4h without rotation. Samples were then brought to a 120 μL volume with nuclease-free water and DNA was sheared using a Covaris S220 instrument (2 cycles, each 50 sec long; 10% duty; 4 intensity; 200 cycles/burst). Dynabeads MyOne Streptavidin C1 beads (Thermo Fisher Scientific) were used to pull down biotinylated fragments according to manufacturer’s guidelines (https://tools.thermofisher.com/content/sfs/manuals/dynabeads_myone_savC1_man.pdf). Libraries bound to Dynabeads MyOne Streptavidin C1 beads were end repaired using the NEBNext Ultra DNA Library Prep Kit for Illumina (New England Biolabs). The beads were then separated on a magnetic stand and washed twice using 1xB&W + 0.1% Triton X-100 and transferred to a 1.5 mL tube. A final wash with 10 mM Tris pH 8.0 was performed and beads were resuspended in 50 μL of the same solution. Final amplification was done in 4 parallel reactions per sample as follows: 10 μL of the library bound to the beads, 25 μL of 2x NEBNext Ultra II Q5 Master Mix, 5 μL of 10 μM Universal PCR primer, 5 μL of 10 μM Indexed PCR primer and 10 μL of nuclease-free water. The number of cycles needed ranged from 10-12 PCR cycles (10 cycles: GFPp_1, GFPp_2 and GFPn_2; 12 cycles: GFPn_1 sample). Each sample was individually barcoded as follows: #2 (GFPp_2), #4 (GFPn_1 and GFPn_2) and #6 (GFPp_1). The PCR reactions were run using the following program: 98°C for 1 min, (98°C for 10 s, 65°C for 75 s, ramping 1.50°C/s) repeated 10-12 times, 65°C for 5 min, 4°C hold. After the amplification, the four reactions were combined into one tube (up to 110 μL volume with nuclease-free water) and then samples were size-selected using Ampure XP beads (Beckman Coulter) following manufacturer’s guidelines (https://genome.med.harvard.edu/documents/sequencing/

Agencourt_AMPure_Protocol.pdf). Following separation, the supernatant contained the final in situ Hi-C library. Libraries were quantified using Qubit dsDNA HS assay kit on a Qubit (Thermo Fisher Scientific) and using an Agilent DNA 1000 kit on a 2100 Bioanalyzer instrument (Agilent). Samples were first pooled and sequenced on an Illumina MiSeq (2×84bp paired-end; MiSeq reagent kit v3-150cycles) to assess library quality. Once the libraries were analysed and past the quality control, they were sequenced on an Illumina NextSeq (2×80bp paired-end; NextSeq 500/550 High Ouput kit v2-150 cycles).

### Quantification and Statistical Analysis

#### Hi-C data processing

For all libraries, FASTQ files were mapped independently, in an iterative fashion against the mm10 reference genome (UCSC) using Bowtie2 with the ‘--very-sensitive’ preset. Briefly, unmapped reads were truncated by 8bp and realigned iteratively, until a valid alignment could be found or the truncated read was shorter than 30bp. Only uniquely mapping reads with a mapping quality (MAPQ) >= 30 were retained for downstream analysis. Restriction fragments were computationally predicted using the Biopython “Restriction” module.

Reads were assigned to fragments, and fragments pairs were formed according to read pairs. Pairs were then filtered for self-ligated fragments, PCR duplicates (both read pairs mapping within 2bp of each other), read pairs mapping further that 5kb from the nearest restriction sites, and ligation products indicating uninformative ligation products (Cournac et al. 2012). The Hi-C matrix was built by binning a genome at a given resolution (e.g. 25kb) and counting valid fragment pairs falling into each respective pair of bins. Finally, bins were masked that have less than 10% of the median number of fragments per bin, and the matrix was normalized using KR matrix balancing (Knight and Ruiz 2013) on each chromosome independently.

#### Observed/expected (OE) Hi-C matrix generation

For each chromosome, we obtained the expected Hi-C contact values by calculating the average contact intensity for all loci at a certain distance. We then transformed the normalized Hi-C matrix into an observed/expected (O/E) matrix by dividing each normalized observed by its corresponding expected value.

#### A-B compartment profiles

We followed a previously described procedure for A-B compartment calculation with minor modifications (Flyamer et al. 2017). Briefly, we transformed the OE matrix for each chromosome at 500kb resolution into a correlation matrix by calculating the correlation of row i and column j for each (i, j). We then calculated the first eigenvector of this matrix, in which positive values indicate the A, and negative values the B compartment, respectively. Sometimes the eigenvector entry signs can be inverted – to ensure that we are assigning the correct sign to individual regions, we also calculated the GC content of each region (correlated with A-B compartment calls (Imakaev et al. 2012; Lieberman-Aiden et al. 2009) and inverted the eigenvector sign if the average GC content of negative-eigenvector entries is higher than that of positive-eigenvector entries. Regions were then sorted according to eigenvector entries, and grouped by percentile (group size here 2%). For each pair of groups, we calculated the average OE values of all corresponding region pairs, which was plotted in the compartment profile. For the calculation of compartment strength, we generated a compartment profile on 1Mb matrices with a group size of 20%. The compartment strength was then calculated as (AA+BB)/AB2 as done previously (Flyamer et al. 2017).

For enrichment of structural changes in A-B compartments the procedure above was followed for matrices at 100kb resolution in mESC Hi-C maps. Regions with positive eigenvector entries were assigned the A, regions with negative eigenvector entries the B compartment label.

#### Average Hi-C feature analysis (TADs, loops, boundaries)

In general, average feature analysis was performed by extracting subsets of the OE matrix (these can be either single regions along the diagonal, or region pairs corresponding the matrix segments off the diagonal) and averaging all resulting sub-matrices. If the sub-matrices were of different size, they were interpolated to a fixed size using “imresize” with the “nearest” setting from the Scipy Python package. TADs and loops annotated in CH12.LX were obtained from (Rao et al. 2014) and lifted over to the mm10 genome version using the UCSC genome browser liftOver tool. The region size for TADs was chosen as 3x TAD size, centred on the TAD, and aggregate analyses were performed in 25kb matrices. The region size around loop anchors was chosen as 300kb in 10kb matrices. Aggregate analyses of OE matrices at MERVL were performed in 25kb matrices in a 500kb window around the central region.

TAD strength was calculated as in (Flyamer et al., 2017). Briefly, we calculated the sum of values in the OE matrix in the TAD-region and the sum of values for the two neighbouring regions of the same size divided by two. The TAD strength was then calculated as the ratio of both numbers. Loop strength was calculated as in (Flyamer et al., 2017). Briefly, we first calculated the sum of all values in the 300kb region of the Hi-C matrix centred on the loop anchors. As a comparison, we calculated the same value for two control regions, substituting one of the loop anchors for an equidistant region in the opposite direction. The loop strength was then calculated as the original sum of values divided by the average sum of values in the two control regions. Hi-C “de novo boundary” aggregate plots are centred on 5’ to 3’-oriented MERVL and show a window of 500kb around the element.

#### Insulation score

We calculated the insulation score as originally defined (Crane et al. 2015) with minor modifications. Briefly, for each region i in the genome, we calculated the average number of interactions in a quadratic window with the lower left corner at (i−1, i+1), and the top right corner at (i−d, i+d), where d is the window size in bins (window size in base pairs can be obtained by multiplying d with the resolution / bin size of the respective matrix). We normalised insulation scores by dividing each region’s score by the average scores of the nearest 300 regions, and log2-transforming the resulting vector, thus accounting for local biases in insulation.

#### Identification of structural changes

To identify genomic regions that undergo structural changes upon the mESC to 2C-like transition, we first calculated the difference between insulation scores in 2CLC and mESC cells in Hi-C matrices of 10kb resolution with three different window sizes: 70kb, 100kb, and 200kb. We then called boundaries on the difference vector as previously described (Crane et al. 2015; Hug et al. 2017), assigning a score to each local minimum corresponding to its depth (i.e. the strength of the insulation change). For stringency, we chose a different cutoff for the boundary score at each window size (0.3, 0.2, and 0.1, respectively), and only label a boundary region as “structural change” if all scores pass the cutoffs.

#### MERVL and other transposable elements in the C57BL/6J reference genome

Repetitive element locations were downloaded from the RepeatMasker website (versions: mm10=4.0.5). Elements with the same RepeatMasker ID were merged. MERVL elements in particular were classified as “complete” if its internal part (MERVL-int) was flanked by two MT2_MM elements facing in the same direction, separated by less than 7kb. Remaining MT2_MM elements are classified as “solo LTR”. Instances of isolated MERVL-int elements were not considered in this analysis.

#### Mappability calculation

Mappability of each base in the mm10 reference genome was calculated by tiling the genome into 36bp long reads and mapping them back to the reference using Bowtie (1.1.1) with default parameters.

#### Division of MERVL by Dux binding

To determine whether the observed chromatin remodelling was dependent on Dux binding to MERVL elements, we used HA-tagged Dux ChIP-seq signal in 2CL cells, induced via Dux overexpression (Hendrickson et al. 2017), to subdivide complete MERVL and solo LTRs into those bound by Dux (Dux+), and not bound by Dux (Dux-) (Figure 2E). A small number of elements in each group have very low mappability (average mappability < 50% in a 100bp region overlapping the 5’ end of the element), so that Dux binding could not be determined (unmappable).

#### Enrichment of features at structural changes

To quantify the association of different genomic features (up-/down-regulated genes, reprogramming-resistant regions, A/B compartments, transposable element types), we first determined the number of features belonging to a group that fall within 30kb of a region with structural changes (n). We then randomised feature locations on each chromosome separately and calculated the number of randomly located features falling within 30kb of a region with structural changes (n_r_i_). The latter was repeated 1000 times to obtain a distribution of the expected number features near regions of structural change. Finally, the Z-score used to quantify the association of features and structural changes is calculated as (n – mean(n_r_i_)) / std(n_r_i_).

#### RNA-seq, ChIP-seq, and ATAC-seq data processing

RNA-seq data were trimmed by quality using Sickle when necessary. Reads were mapped to the mouse reference genome (v GRCm38 downloaded from the Illumina iGenome’s portal [http://support.illumina.com/sequencing/sequencing_software/igenome.ilmn]) using TopHat2 with default parameters, taking into consideration the minimum and maximum intron size of the mouse genome (-g 2 -i 30 -I 1050000 --min-segment-intron=30 --max-segment-intron=1050000). Uniquely mapping reads were extracted using the NH:i flag reported by TopHat2. Normalized coverage BigWig tracks for RNA-seq data were generated from the resulting BAM files using bamCoverage from the deepTools package with a window size of 500bp, and normalization by RPKM. Lists of differentially expressed genes were obtained from (Ishiuchi et al. 2015).

ChIP-seq data were mapped to the same genome version using Bowtie2. Uniquely mapping reads were extracted using the XS:i flag reported by Bowtie2. For samples with input or equivalent control, MACS2 was used to calculate the fold enrichment BigWig tracks used for plotting.

Adapters were trimmed from ATAC-seq reads using BBDuk. Reads were mapped to the mm10 reference genome with Bowtie2 and filtered for multimapping reads as with ChIP-Seq data. Further filtering of reads mapping to the mitochondrial genome was performed and PCR duplicates were removed with Picard tools. Coverage of uniquely mapping reads was normalized by number of mapped reads and converted to BigWig format using pybedtools.

#### Dux ChIP-seq peaks and MERVL binding

Reads from the Dux-HA expression ChIP-seq dataset were mapped and filtered as described in the previous section. Peaks were called with the callpeak command of MACS2 with the paired-end bam (BAMPE) format. Bedtools intersect was used on the narrowPeak output of MACS2 and the MERVL annotation (for complete and solo LTRs, respectively) to separate Dux-bound from not-bound copies.

#### Enrichment heatmaps and meta-profiles for genomic datasets

Each row in a heatmap corresponds to a specific genomic region. Plotted in each row is the binned signal of one of the processed datasets above (RNA-seq, ChIP-seq, ATAC-seq), surrounding that region. To obtain the signal for each row, intervals and scores from processed BigWig tracks are extracted in a window of fixed size centred at the element with a 5’ to 3’-end orientation. Scores were binned into 100 bins, using a mean weighted by the overlap of each interval with the respective bin. Meta-profiles were obtained by calculating a 5% trimmed mean for each column of the heatmap. For visualization (but not for meta-profile calculation), rows with more than 50% of NaN values or unmappable bins are omitted. Difference heatmaps between two types of cells or stages have been obtained by simply subtracting enrichment heatmaps for a specific dataset from one another.

#### Insulation change and expression change correlation analysis

We obtained difference heatmaps (250kb window) for expression and insulation data from mESC minus 2CLC, centred at elements of interest (complete MERVL, solo LTR, IAPEZ, L1_MM, Dux peaks without MERVL). For each row, we calculated the correlation of expression changes with inverse insulation changes for all bins downstream of the element. Correlation values were then visualized in a boxplot. We proceeded similarly for *in vivo* expression and insulation difference heatmaps in the early 2-cell minus 8-cell stages.

#### MERVL long-range contact enrichment

We selected all pairs of MERVL separated by a distance of less than 5Mb and used these as input for an average Hi-C feature analysis (see above). We calculated aggregate matrices for MERVL in mESC and 2CLC. In addition, we generated a set of control regions by substituting one MERVL in each pair by an equidistant region in the opposite direction. Statistical testing was performed on O/E values between pairs of MERVL, using mESC and 2CLC results as the two groups in a Wilcoxon signed-rank test.

#### MERVL directionality analysis

Directionality bias for MERVL was assessed as previously described for CTCF (Rowley et al., 2017). Briefly, the analysis centres on the 5’ end of complete MERVL and computes the log2-ratio of Hi-C contact values to all bins up- and downstream of the element, as a function of distance.

#### Motif enrichment

The callpeak command from MACS2 was run on the in-vitro (Hendrickson et al., 2017) and in vivo (Wu et al., 2016) ATAC-seq data. For the in vivo data, we calculated the difference in insulation scores between mESC and 2CLC, identifying regions with a change larger than 0.2. A 40kb window surrounding these regions was intersected with the in vitro ATAC-seq peaks and used for motif analysis. For the in vitro data, we used 40kb windows surrounding complete MERVL annotated in C57BL/6J, intersected with the in vivo ATAC-seq peaks. Motif search was run using Homer findMotifsGenome.pl on the mm10 reference genome with default parameters, identifying both known and novel DNA motifs.

#### Chimeric read analysis to detect MERVL polymorphisms

To find strain-specific differences in MERVL integration compared to the C57Bl/6J reference genome, we utilised aligned reads from DNA sequencing data generated in the alternative strain. In principle, the type of data (Hi-C, ATAC-seq, ChIP-seq) is irrelevant, although some approaches might have a higher density of reads in certain genomes, depending on the biological question or experimental setup, thus increasing the likelihood of identifying polymorphisms in those regions. We use DNA sequencing reads to (i) confirm the presence MERVL annotated in C57BL/6J in the alternative strain, (ii) confirm the absence of MERVL annotated in C57BL/6J in the alternative strain, (iii) find novel integrations of MERVL/MT2_MM in the alternative strain.

(i) For each complete MERVL annotated in C57BL/6J, we used samtools on the aligned read BAM files to extract all uniquely mapping reads overlapping either the 5’ or the 3’ end. We then extracted the longest consecutive mapping portion of the read using the read’s CIGAR string (largest “M” entry). We called a MERVL confirmed if at least 10 reads overlap either the 5’ or 3’ region of the element by more than 10bp. We also used this approach on the sperm Hi-C datasets from (Du et al. 2017) and (Ke et al. 2017) to obtain a high-confidence set of MERVL shared between C57BL/6J and PWK/PhJ, and DBA/2j, respectively.

(ii) We mapped the sequencing reads with BWA-mem (default settings). Paired-end reads were mapped as two single-end datasets. We then extracted all reads that are part of a chimeric pair where both reads map in the same orientation on the same chromosome less than 10kb apart. For Hi-C data, we first ensured that the chimeric pair is not the result of a Hi-C ligation by excluding read pairs that form a junction motif of the respective restriction enzyme. We considered the region spanned by a chimeric read pair likely to be missing from the alternative strain. A MERVL annotated in C57BL/6J is called “missing” from the alternative genome, if a minimum of 10 missing genomic regions span at least 99% of the element, thus reducing false-positives due to misaligned reads.

(iii) We proceeded with the mapping of reads as in (ii). We then extracted all chimeric read pairs (excluding Hi-C ligation junctions as in (ii)) where one read is uniquely mapping and the other one maps to multiple locations in the genome. If the multi-mapping half maps to any of the MT2_MM elements in the reference genome, the uniquely mapping half likely represents the location of an unannotated MERVL insertion site in the alternative strain. We called a novel integration site if at least 10 such chimeric read pairs could be found at this exact location. Finally, we removed all identified integration sites that overlap annotate MERVL or MT2_MM as false-positives.

#### Virtual 4C to identify eGFP integration site

To detect the insertion site(s) of the MERVL-driven eGFP reporter used to identify 2CLC, we map our Hi-C sequencing reads against a Bowtie2 index that also contains the sequence of the pEGFP N2 vector (Clontech) used in our experiments (for this, we removed the CMV promoter sequence, as it has been replaced with a MERVL LTR). Virtual 4C was performed by plotting all Hi-C contacts the (truncated) pEGFP sequence makes throughout the genome. We identified the integration site as the region with the highest contact intensity throughout the genome.

#### List of experimental material

| Reagent | Source | Identifier |
| --- | --- | --- |
| 3 M Sodium acetate buffer (pH 5.2) | Sigma-Aldrich | S7899-100ml |
| 5 U/ μL DNA plimerase I klenow Fragment | New England Biolabs | M0210L |
| agarose basic applichem | Applichem | A8963,1000 |
| Agencourt AMPure XP beads | Beckman Coulter | A63881 |
| Agilent DNA 1000 Just Reagents | Agilent | 5067-1505 |
| Agilent DNA 1000 Kit | Agilent | 5067-1504 |
| Biotin-14-dCTP | Thermo Fisher Scientific | 19518-018 |
| Blunt/TA Ligase Master Mix | New England Biolabs | M0367L |
| BSA-Molecular Biology Grade (20 mg/ml) | New England Biolabs | B9000S |
| cOmplete ULTRA Tablets, Mini. EASY Pack | Roche (Sigma-Aldrich) | 05 892 970 001 |
| dATP | Thermo Fisher Scientific | R0141 |
| dCTP | Thermo Fisher Scientific | R0151 |
| dGTP | Thermo Fisher Scientific | R0161 |
| DTT=1-4-Dithiothreitol | Roth | 6908.1 |
| dTTP | Thermo Fisher Scientific | R0171 |
| Dynabeads® MyOne™ Streptavidin C1 | Thermo Fisher Scientific | 65001 |
| EDTA solution pH 8.0 (0.5 M) Molecular biology | Applichem | A4892,0500 |
| Ethanol absolute (>99.8%) | Sigma-Aldrich | 24102-1L-R |
| Formaldehyde Sol. 37% for molecular biology | VWR International Gmb | 437536C |
| Glacial Acetic Acid | AMRESCO | 0714-500 mL |
| Glycine Molecular Biology grade | Applichem | Applichem |
| GlycoBlue (15 mg/ml) (0.3 ml Tube) | Thermo Fisher Scientific | AM9516 |
| High Sensitivity DNA Analysis Just Reagents | Agilent | 5067-4627 |
| High Sensitivity DNA Analysis Kits | Agilent | 5067-4626 |
| IGEPAL CA-630 | Sigma-Aldrich | 18896-100ml |
| Magnesium chloride | Santa Cruz Biotechnology, Inc | 7786-30-3 |
| Mbol | New England Biolabs | R0147L |
| Phenol - chloroform - isoamyl alcohol mixture, BioUltra, for molecular biology, 25:24:1 | Sigma-Aldrich | 77617-100ML |
| Proteinase K solution | Applichem-Panreac | A4392,0005 |
| Qubit® Assay Tubes | Thermo Fisher Scientific | Q32856 |
| Qubit® dsDNA HS Assay Kit | Thermo Fisher Scientific | Q32854 |
| RNAse A | Applichem | A3832,0050 |
| SDS Molecular biology grade | Applichem | A2263,0100 |
| Sodium chloride | CALBIOCHEM | 567440 |
| T4 DNA Ligase | Thermo Fisher Scientific | EL0012 |
| T4 DNA Ligase Reaction Buffer (10X) | Thermo Fisher Scientific | B69 |
| T4 DNA polymerase | New England Biolabs | M0203L |
| Tris Base = Tris Molecular biology grade (MW=121.1) | Applichem | A2264,1000 |
| Tris buffer Molecular biology grade pH 7.4 (1M) | Applichem | A3981,1000 |
| Tris buffer Molecular biology grade pH 8.0 (1M) | Applichem | A4577,1000 |
| Tris buffer pH 8.0 (1 M) Molecular biology grade | Applichem | A4577,1000 |
| Tris buffer pH 8.0 (1 M) Molecular biology grade | Applichem | A4975,0500 |
| NEBNext® End Repair Module | New England Biolabs | E6050L |
| NEBNext® Multiplex Oligos for Illumina® (Index Primers Set 1) | New England Biolabs | E7335L |
| NEBNext® Q5® Hot Start HiFi PCR Master Mix | New England Biolabs | M0543L |
| NEBNext® ltra™ DNA Library Prep Kit for Illumina® | New England Biolabs | E7370L |
| NEBNext® Ultra™ End Repair/dA-Tailing Module | New England Biolabs | E7442L |

#### List of applied software

| Name (version) | Publication / authors | Website |
| --- | --- | --- |
| Bowtie2 (v2.2.4) | (Langmead and Salzberg 2012) | http://bowtie-bio.sourceforge.net/bowtie2/index.shtml |
| BWA (v0.7.17) | (Li and Durbin 2009) | http://bio-bwa.sourceforge.net/ |
| MACS2 (v2.1.1) | (Zhang et al. 2008) | https://github.com/taoliu/MACS |
| Homer (v4.8) | (Heinz et al. 2010) | http://homer.ucsd.edu/homer/ |
| HiC-Pro (v2.9.0) | (Servant et al. 2015) | https://github.com/nservant/HiC-Pro |
| Python (v2.7.12 and v3.6.4) | Python Software Foundation | https://www.python.org |
| Biopython | Biopython Contributors | http://biopython.org/ |
| Bedtools (v2.26.0) | (Quinlan and Hall 2010) | http://bedtools.readthedocs.io/ |
| Samtools (v1.6) | (Li et al. 2009) | http://www.htslib.org/ |
| Scipy (v0.19.0) | Scipy developers | https://www.scipy.org/ |
| deepTools (v3.0.1) | (Ramírez et al. 2016) | https://deeptools.readthedocs.io |
| BBMap (v36.88) | JGI | https://jgi.doe.gov/data-and-tools/bbtools/ |
| Sickle (v1.33) | Joshi and Fass | https://github.com/najoshi/sickle |
| Picard-tools (v1.106) | Broad Institute | https://broadinstitute.github.io/picard/ |
| SNPsplit (v0.3.2) | (Krueger and Andrews 2016) | https://www.bioinformatics.babraham.ac.uk/projects/SNPsplit/ |
| IGV (v2.3.88) | (Robinson et al. 2011) | http://software.broadinstitute.org/software/igv/ |
| TopHat2 (v2.1.0) | (Kim et al. 2013) | https://ccb.jhu.edu/software/tophat/index.shtml |
| Stringtie (v.1.3.1) | (Pertea et al. 2015) | https://ccb.jhu.edu/software/stringtie/ |
| liftOver | (Kent et al. 2002) | http://hgdownload.soe.ucsc.edu/admin/exe/linux.x86_64/ |

## EXTENDED DATA FIGURES

**Extended Data Figure 1.**
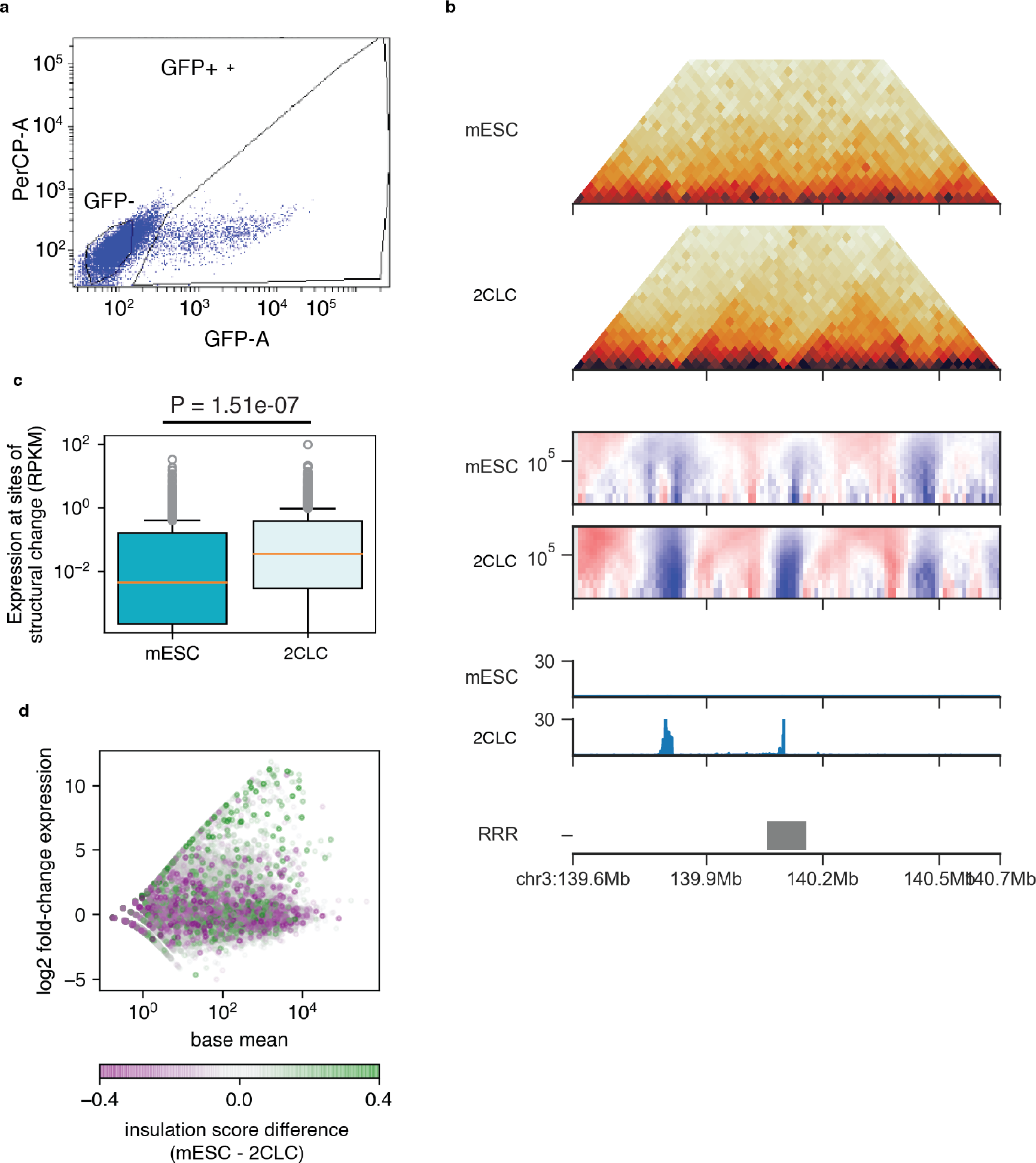
FACS sorting of mESC and 2CLC and analysis of expression changes from 2CLC and mESC. (a) FACS gating plot for sorting GFP+ (2CLC) and GFP−(mESC) cells. (b) Representative example region on chromosome 2 highlighting changes in Hi-C, insulation, and expression in 2CLC compared to mESCs at reprogramming resistant regions (RRR). (c) Boxplots showing the distribution of expression (RPKM) values at regions undergoing structural changes in 2CLC. (d) MA plot showing changes in gene expression between 2CLC and mESC. Points correspond to genes and are color-coded by the magnitude of structural change in 2CLC.

**Extended Data Figure 2.**
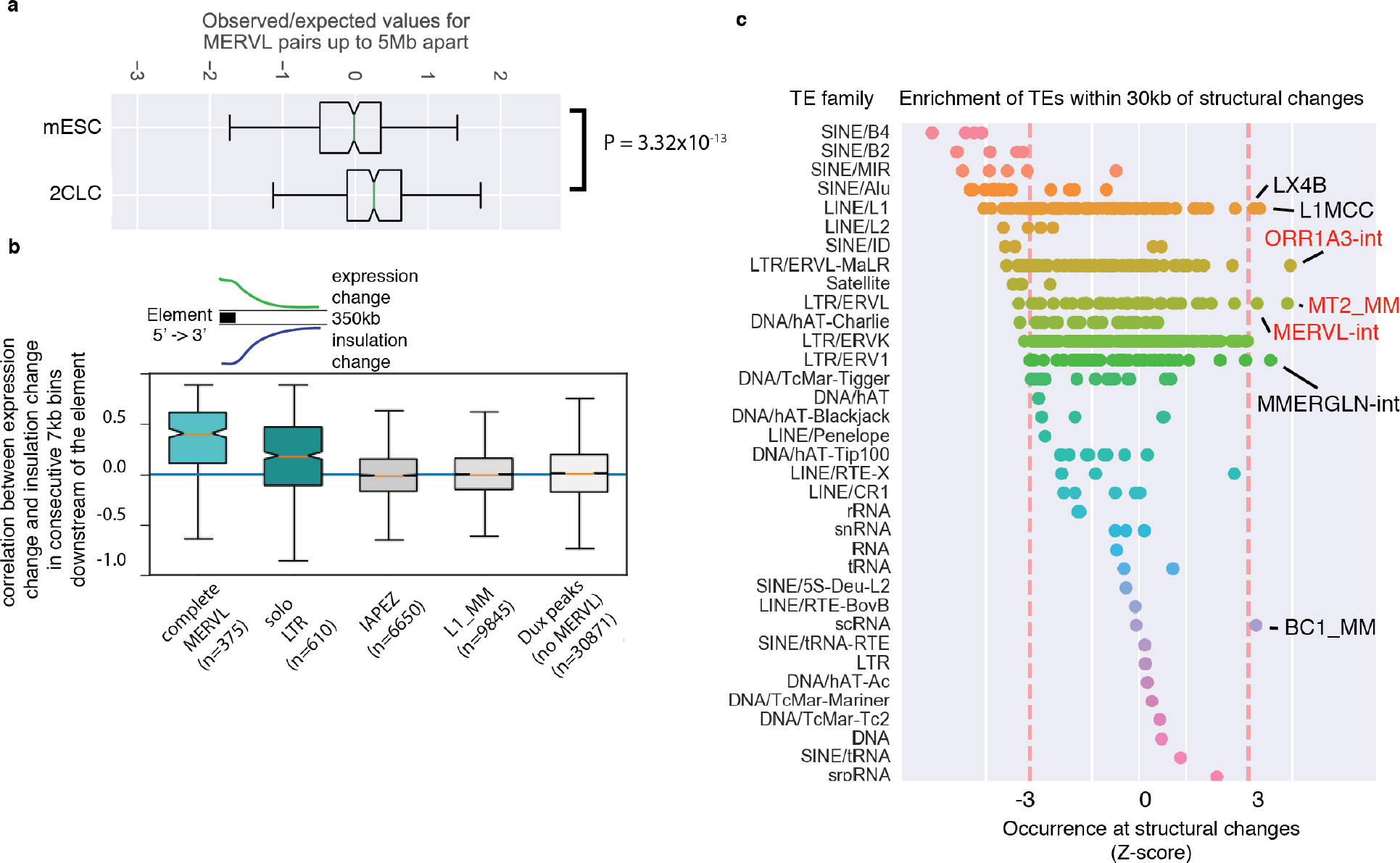
Features associated with novel insulating regions in 2CLC. (a) Boxplot of the observed vs expected Hi-C interaction values for complete MERVL up to 5Mb apart in mESCs and 2CLC. (b) Boxplots of expression change vs insulation change correlations in regions downstream of individual elements (complete MERVL, solo LTRs, IAPEZ, L1_MM, and Dux peaks without MERVL). (c) Association of types of transposable elements with structural changes. X axis shows all transposable element families. Points denote TE types. Y axis denotes Z-score quantifying the likelihood of a TE type to occur at regions of structural change (higher values = more likely, lower values = less likely). TE types with a Z-score > 3 are explicitly labelled, red labels denote components of MERVL.

**Extended Data Figure 3.**
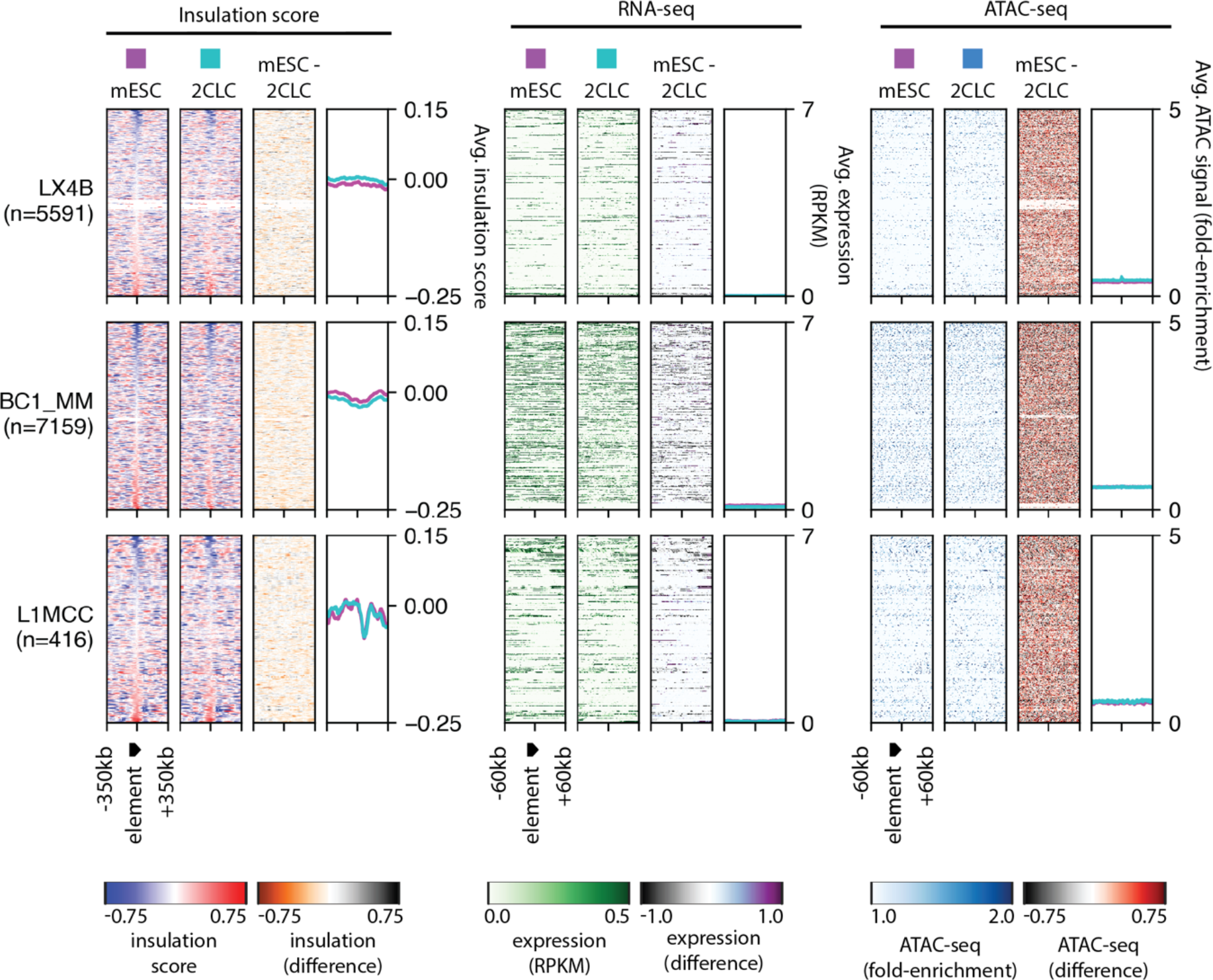
Other types of TEs do not display coordinated insulation, expression, or chromatin opening changes as a whole. Insulation score, expression, and chromatin openness data at all members of LX4B, BC1_MM, and L1MCC

**Extended Data Figure 4.**
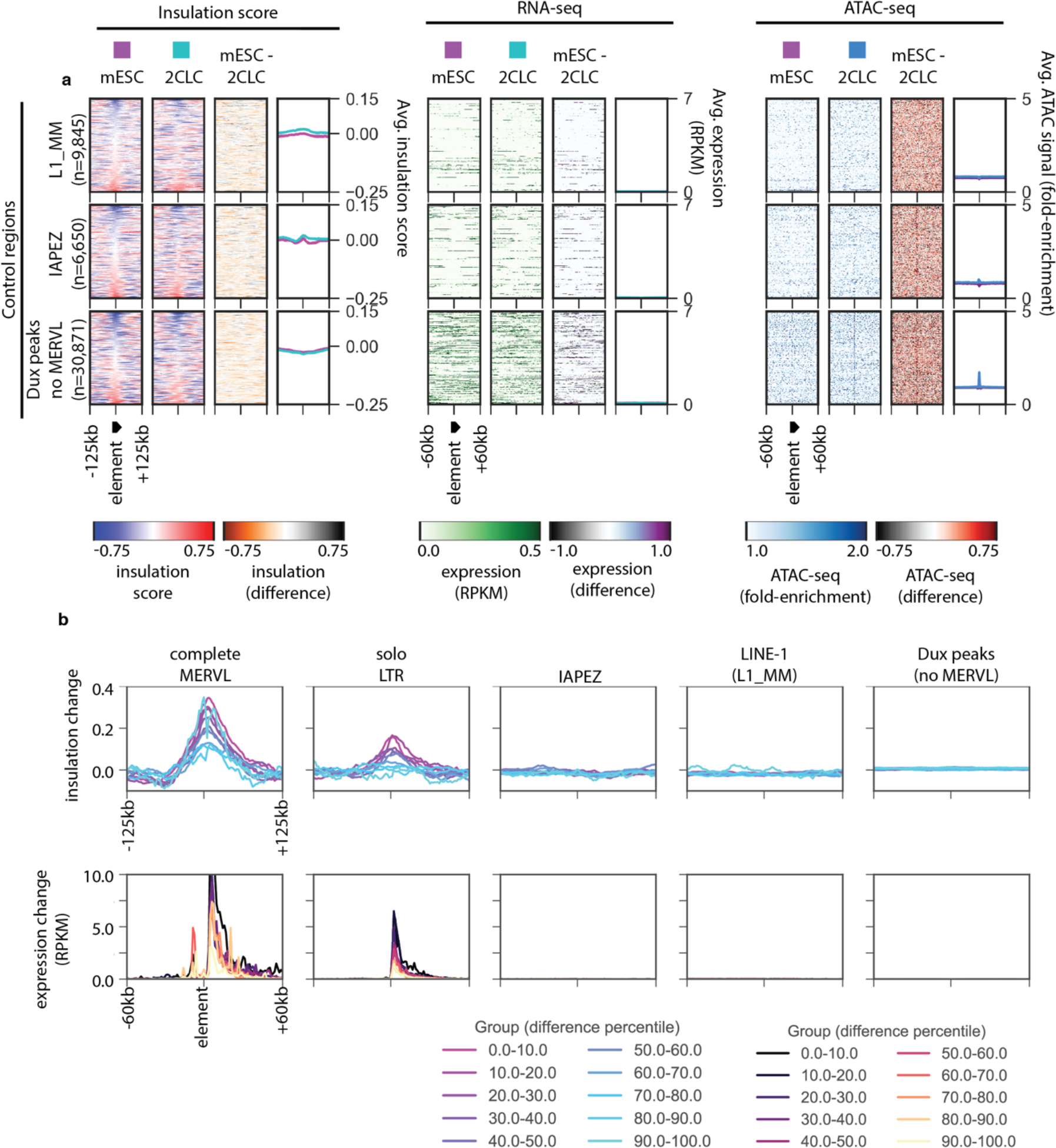
Examination of chromatin and expression changes at various control regions. (a) Insulation score, expression, and chromatin openness data at L1_MM and IAPEZ elements, as well as HA-tagged Dux ChIP-seq peaks that do not overlap with MERVL. (b) Metaplots of insulation (top) and expression (bottom) changes from mESC to 2CLC. Changes are stratified by percentiles of the magnitude of change, i.e. the regions with the largest insulation changes are in the 0-10th percentile, and so on.

**Extended Data Figure 5.**
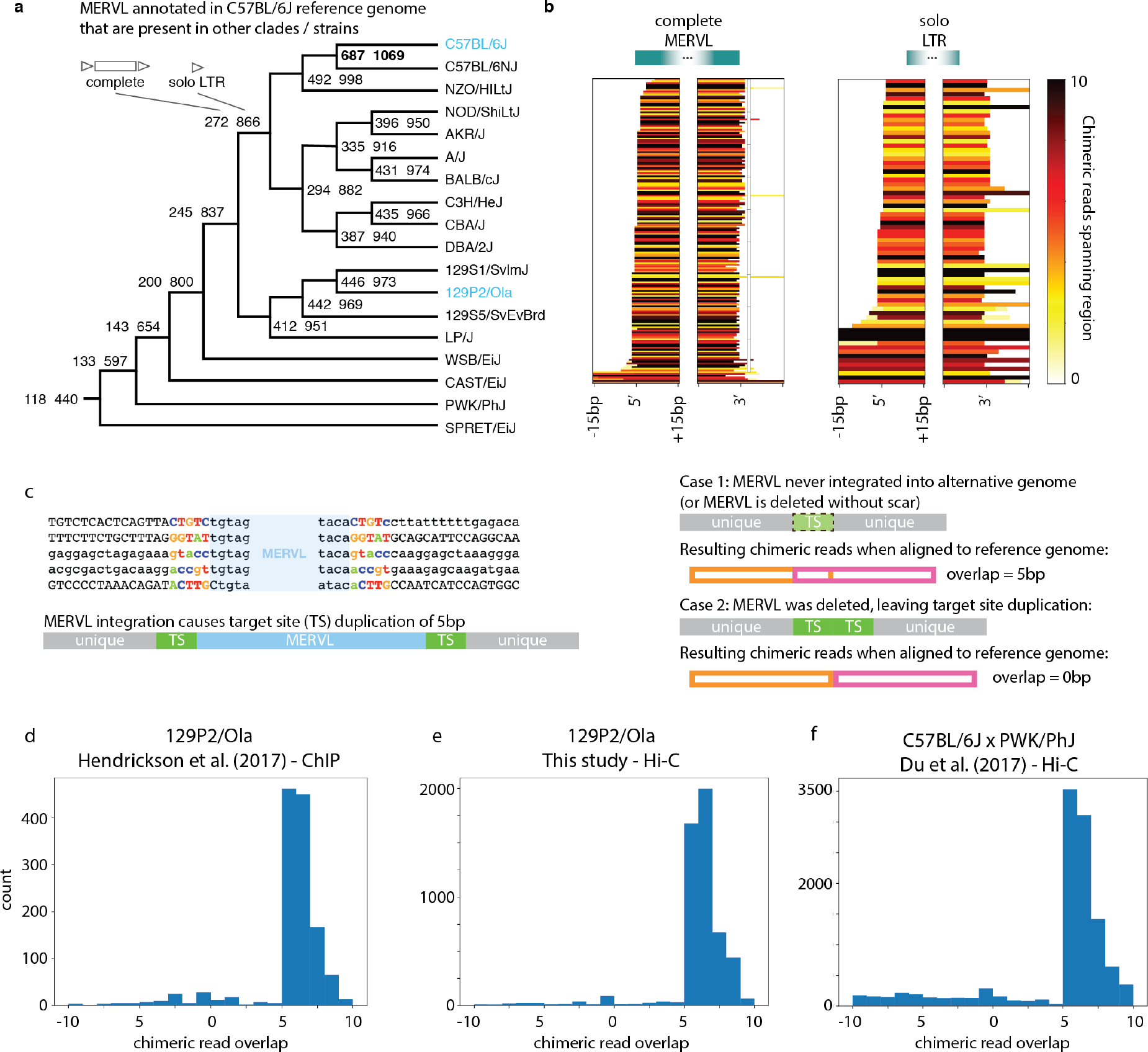
MERVL polymorphism in 18 different mouse strains. (a) Phylogenetic tree of 18 mouse strains annotated with the number of MERVL (annotated in C57BL/6J) that are common between strains/clades. (b) Heatmap of gaps between chimeric reads overlapping complete MERVL and solo LTRs. (c) Example of 5bp target site duplication (TSD) at MERVL in C57BL/6J and schematic of MERVL integration and subsequent deletion or non-integration for chimeric reads at the site of the annotated MERVL. In case 1 (non-integration), MERVL never integrated into the alternative genome and hence did not duplicate the 5bp target site. Corresponding chimeric reads “share” the 5bp target site, and hence overlap by approximately 5bp. In case 2 (deletion of MERVL leaving TSD), the target site is duplicated and would therefore appear twice in the chimeric read(s). There would be no overlap between the two halves of the chimeric read. (d-f) Distributions of the overlap between chimeric read halves in (d, e) 129P2/Ola, and (f) PWK/PhJ. Peaks at 5 indicate TSDs are not present in either strain.

**Extended Data Figure 6.**
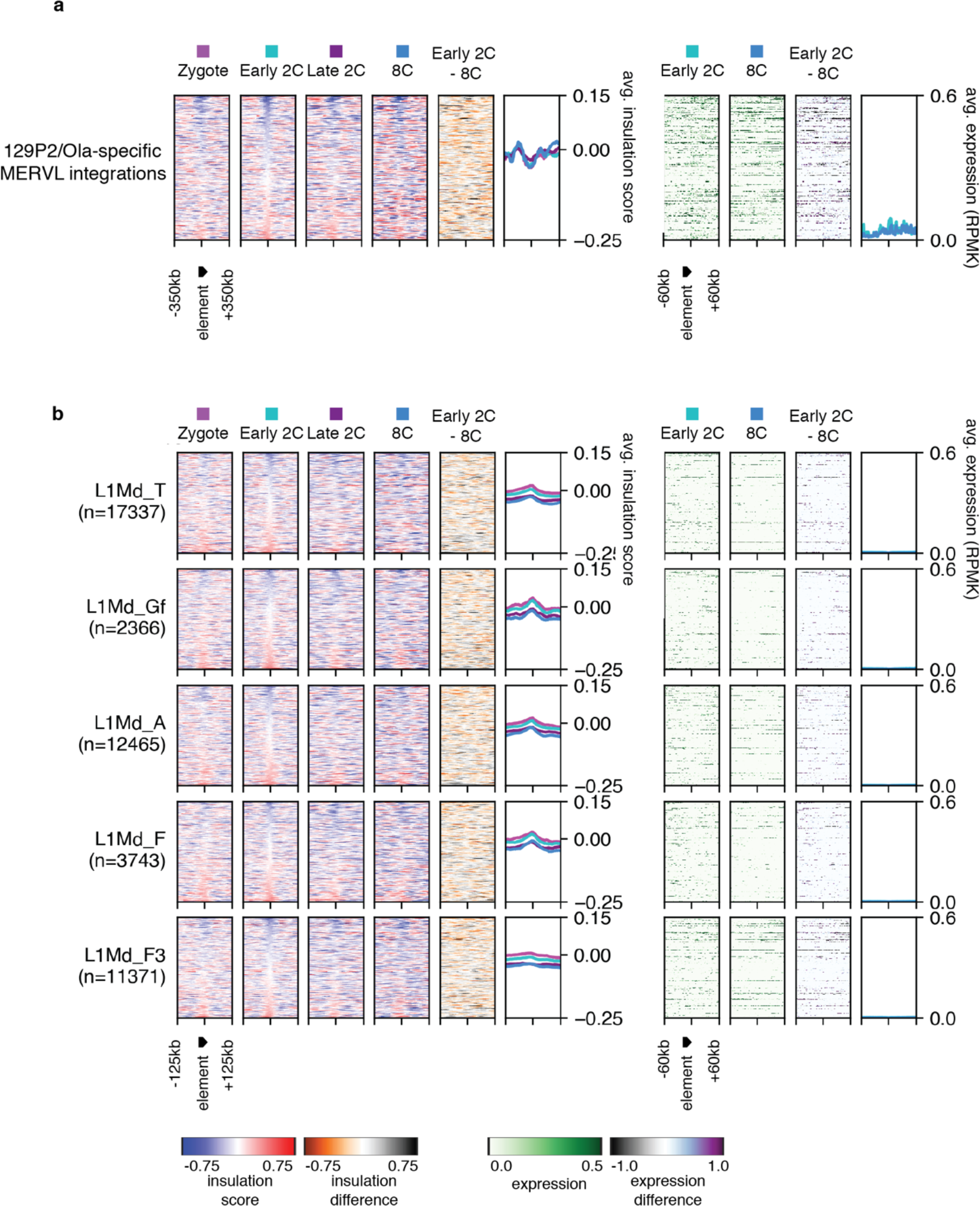
Heatmaps of insulation score and expression for (a) 129P2/Ola-specific MERVL integrations in a C57BL/6J x PWK-PhJ background, and (b) different types of LINE1 elements throughout early embryonic development.

## SUPPLEMENTARY TABLES

**Supplementary Table 1. Low input Hi-C statistics for mESC and 2CLC.** Table provided as an Excel file (Kruse_et_al-Supplementary-Table-1.xlsx).

**Supplementary Table 2. List of regions in 2CLC that gain insulation compared to mESC.** Table provided as an Excel file (Kruse_et_al-Supplementary-Table-2.xlsx).

## REFERENCES

Ancelin K, Syx L, Borensztein M, Ranisavljevic N, Vassilev I, Briseño-Roa L, Liu T, Metzger E, Servant N, Barillot E, et al. 2016. Maternal LSD1/KDM1A is an essential regulator of chromatin and transcription landscapes during zygotic genome activation. Elife 5: e08851.

Bonev B, Mendelson Cohen N, Szabo Q, Fritsch L, Papadopoulos GL, Lubling Y, Xu X, Lv X, Hugnot JP, Tanay A, et al. 2017. Multiscale 3D Genome Rewiring during Mouse Neural Development. Cell 171: 557–572.e24.

Bošković A, Eid A, Pontabry J, Ishiuchi T, Spiegelhalter C, Raghu Ram EVS, Meshorer E, Torres-Padilla ME. 2014. Higher chromatin mobility supports totipotency and precedes pluripotency in vivo. Genes Dev 28: 1042–1047.

Bourque G, Leong B, Vega VB, Chen X, Yen LL, Srinivasan KG, Chew JL, Ruan Y, Wei CL, Huck HN, et al. 2008. Evolution of the mammalian transcription factor binding repertoire via transposable elements. Genome Res 18: 1752–1762.

Burton A, Torres-Padilla ME. 2014. Chromatin dynamics in the regulation of cell fate allocation during early embryogenesis. Nat Rev Mol Cell Biol 15: 722–734.

Copeland NG, Hutchison KW, Jenkins NA. 1983. Excision of the DBA ecotropic provirus in dilute coat-color revertants of mice occurs by homologous recombination involving the viral LTRs. Cell 33: 379–387.

Cournac A, Koszul R, Mozziconacci J. 2016. The 3D folding of metazoan genomes correlates with the association of similar repetitive elements. Nucleic Acids Res 44: 245–255.

Crane E, Bian Q, McCord RP, Lajoie BR, Wheeler BS, Ralston EJ, Uzawa S, Dekker J, Meyer BJ. 2015. Condensin-driven remodelling of X chromosome topology during dosage compensation. Nature 523: 240–244.

Dahl JA, Jung I, Aanes H, Greggains GD, Manaf A, Lerdrup M, Li G, Kuan S, Li B, Lee AY, et al. 2016. Broad histone H3K4me3 domains in mouse oocytes modulate maternal-to-zygotic transition. Nature 537: 548–552.

Darrow EM, Huntley MH, Dudchenko O, Stamenova EK, Durand NC, Sun Z, Huang S-C, Sanborn AL, Machol I, Shamim M, et al. 2016. Deletion of DXZ4 on the human inactive X chromosome alters higher-order genome architecture. Proc Natl Acad Sci 201609643.

De Iaco A, Planet E, Coluccio A, Verp S, Duc J, Trono D. 2017. DUX-family transcription factors regulate zygotic genome activation in placental mammals. Nat Genet 49: 941–945.

de Jong J, Akhtar W, Badhai J, Rust AG, Rad R, Hilkens J, Berns A, van Lohuizen M, Wessels LFA, de Ridder J. 2014. Chromatin landscapes of retroviral and transposon integration profiles. PLoS Genet 10: e1004250.

Díaz N, Kruse K, Erdmann T, Staiger AM, Ott G, Lenz G, Vaquerizas JM. 2018. Chromatin conformation analysis of primary patient tissue using a low input Hi-C method. Nat Commun 9: 4938.

Dixon JR, Selvaraj S, Yue F, Kim A, Li Y, Shen Y, Hu M, Liu JS, Ren B. 2012. Topological domains in mammalian genomes identified by analysis of chromatin interactions. Nature 485: 376–380.

Du Z, Zheng H, Huang B, Ma R, Wu J, Zhang XX, He J, Xiang Y, Wang Q, Li Y, et al. 2017. Allelic reprogramming of 3D chromatin architecture during early mammalian development. Nature 547: 232–235.

Eckersley-Maslin MA, Svensson V, Krueger C, Stubbs TM, Giehr P, Krueger F, Miragaia RJJ, Kyriakopoulos C, Berrens RV V., Milagre III, et al. 2016. MERVL/Zscan4 Network Activation Results in Transient Genome-wide DNA Demethylation of mESCs. Cell Rep 17: 179–192.

Fadloun A, Le Gras S, Jost B, Ziegler-Birling C, Takahashi H, Gorab E, Carninci P, Torres-Padilla ME. 2013. Chromatin signatures and retrotransposon profiling in mouse embryos reveal regulation of LINE-1 by RNA. Nat Struct Mol Biol 20: 332–338.

Feschotte C. 2008. Transposable elements and the evolution of regulatory networks. Nat Rev Genet 9: 397–405.

Friedli M, Trono D. 2015. The Developmental Control of Transposable Elements and the Evolution of Higher Species. Annu Rev Cell Dev Biol 31: 429–51.

Giorgetti L, Lajoie BR, Carter AC, Attia M, Zhan Y, Xu J, Chen CJ, Kaplan N, Chang HY, Heard E, et al. 2016. Structural organization of the inactive X chromosome in the mouse. Nature 535: 575–579.

Hendrickson PG, Doráis JA, Grow EJ, Whiddon JL, Lim J-W, Wike CL, Weaver BD, Pflueger C, Emery BR, Wilcox AL, et al. 2017. Conserved roles of mouse DUX and human DUX4 in activating cleavage-stage genes and MERVL/HERVL retrotransposons. Nat Genet 49: 925–934.

Hug CB, Grimaldi AG, Kruse K, Vaquerizas JM. 2017. Chromatin Architecture Emerges during Zygotic Genome Activation Independent of Transcription. Cell 169: 216–228.e19.

Hug CB, Vaquerizas JM. 2018. The Birth of the 3D Genome during Early Embryonic Development. Trends Genet 34: 903–914.

Ishiuchi T, Enriquez-Gasca R, Mizutani E, Bošković A, Ziegler-Birling C, Rodriguez-Terrones D, Wakayama T, Vaquerizas JM, Torres-Padilla M-E. 2015. Early embryonic-like cells are induced by downregulating replication-dependent chromatin assembly. Nat Struct Mol Biol 22: 662–671.

Jachowicz JW, Bing X, Pontabry J, Bošković A, Rando OJ, Torres-Padilla M-E. 2017. LINE-1 activation after fertilization regulates global chromatin accessibility in the early mouse embryo. Nat Genet 49: 1502–1510.

Ke Y, Xu Y, Chen X, Feng S, Liu Z, Sun Y, Yao X, Li F, Zhu W, Gao L, et al. 2017. 3D Chromatin Structures of Mature Gametes and Structural Reprogramming during Mammalian Embryogenesis. Cell 170: 367–381.e20.

Lieberman-Aiden E, Berkum NL Van, Williams L, Imakaev M, Ragoczy T, Telling A, Amit I, Lajoie BR, Sabo PJ, Dorschner MO, et al. 2009. Comprehensive Mapping of Long-Range Interactions Revelas Folding Principles of the Human Genome. Science 326: 289–93.

Liu X, Wang C, Liu W, Li J, Li C, Kou X, Chen J, Zhao Y, Gao H, Wang H, et al. 2016. Distinct features of H3K4me3 and H3K27me3 chromatin domains in pre-implantation embryos. Nature 537: 558–562.

Macfarlan TS, Gifford WD, Driscoll S, Lettieri K, Rowe HM, Bonanomi D, Firth A, Singer O, Trono D, Pfaff SL. 2012. Embryonic stem cell potency fluctuates with endogenous retrovirus activity. Nature 487: 57–63.

Matoba S, Liu Y, Lu F, Iwabuchi KA, Shen L, Inoue A, Zhang Y. 2014. Embryonic development following somatic cell nuclear transfer impeded by persisting histone methylation. Cell 159: 884–895.

Nellåker C, Keane TM, Yalcin B, Wong K, Agam A, Belgard TG, Flint J, Adams DJ, Frankel WN, Ponting CP. 2012. The genomic landscape shaped by selection on transposable elements across 18 mouse strains. Genome Biol 13.

Peaston AE, Evsikov A V., Graber JH, de Vries WN, Holbrook AE, Solter D, Knowles BB. 2004. Retrotransposons regulate host genes in mouse oocytes and preimplantation embryos. Dev Cell 7: 597–606.

Percharde M, Lin CJ, Yin Y, Guan J, Peixoto GA, Bulut-Karslioglu A, Biechele S, Huang B, Shen X, Ramalho-Santos M. 2018. A LINE1-Nucleolin Partnership Regulates Early Development and ESC Identity. Cell 174: 391–405.e19.

Pope BD, Ryba T, Dileep V, Yue F, Wu W, Denas O, Vera DL, Wang Y, Hansen RS, Canfield TK, et al. 2014. Topologically associating domains are stable units of replication-timing regulation. Nature 515: 402–405.

Rao SSPP, Huntley MH, Durand NC, Stamenova EK, Bochkov ID, Robinson JT, Sanborn AL, Machol I, Omer AD, Lander ES, et al. 2014. A 3D map of the human genome at kilobase resolution reveals principles of chromatin looping. Cell 159: 1665–1680.

Rodriguez-Terrones D, Gaume X, Ishiuchi T, Weiss A, Kopp A, Kruse K, Penning A, Vaquerizas JM, Brino L, Torres-Padilla ME. 2018. A molecular roadmap for the emergence of early-embryonic-like cells in culture. Nat Genet 50: 106–119.

Rodriguez-Terrones D, Torres-Padilla ME. 2018. Nimble and Ready to Mingle: Transposon Outbursts of Early Development. Trends Genet 34: 806–820.

Schmidt D, Schwalie PC, Wilson MD, Ballester B, Goncalves A, Kutter C, Brown GD, Marshall A, Flicek P, Odom DT. 2012. Waves of retrotransposon expansion remodel genome organization and CTCF binding in multiple mammalian lineages. Cell 148: 335–48.

Stadhouders R, Vidal E, Serra F, Di Stefano B, Le Dily F, Quilez J, Gomez A, Collombet S, Berenguer C, Cuartero Y, et al. 2018. Transcription factors orchestrate dynamic interplay between genome topology and gene regulation during cell reprogramming. Nat Genet 50: 238–249.

Sun JH, Zhou L, Emerson DJ, Phyo SA, Titus KR, Gong W, Gilgenast TG, Beagan JA, Davidson BL, Tassone F, et al. 2018. Disease-Associated Short Tandem Repeats Co-localize with Chromatin Domain Boundaries. Cell 175: 224–238.e15.

Thompson PJJ, Macfarlan TSS, Lorincz MCC. 2016. Long Terminal Repeats: From Parasitic Elements to Building Blocks of the Transcriptional Regulatory Repertoire. Mol Cell 62: 766–776.

Wang J, Vicente-García C, Seruggia D, Moltó E, Fernandez-Miñán A, Neto A, Lee E, Gómez-Skarmeta JL, Montoliu L, Lunyak V V, et al. 2015. MIR retrotransposon sequences provide insulators to the human genome. Proc Natl Acad Sci 112: E4428–37.

Winter DJ, Ganley ARD, Young CA, Liachko I, Schardl CL, Dupont P, Berry D, Ram A, Scott B, Cox MP. 2018. Repeat elements organise 3D genome structure and mediate transcription in the filamentous fungus Epichloë festucae. PLoS Genet 14: e1007467.

Wu J, Huang B, Chen H, Yin Q, Liu Y, Xiang Y, Zhang B, Liu B, Wang Q, Xia W, et al. 2016. The landscape of accessible chromatin in mammalian preimplantation embryos. Nature 534: 652–657.

Xu Q, Xie W. 2018. Epigenome in Early Mammalian Development: Inheritance, Reprogramming and Establishment. Trends Cell Biol 28: 237–253.

Zhang B, Zheng H, Huang B, Li W, Xiang Y, Peng X, Ming J, Wu X, Zhang Y, Xu Q, et al. 2016. Allelic reprogramming of the histone modification H3K4me3 in early mammalian development. Nature 537: 553–557.

## REFERENCES

Cournac A, Marie-Nelly H, Marbouty M, Koszul R, Mozziconacci J. 2012. Normalization of a chromosomal contact map. BMC Genomics 13: 436.

Flyamer IM, Gassler J, Imakaev M, Ulyanov S V., Abdennur N, Razin S V., Mirny LA, Tachibana-Konwalski K, Brandão HB, Ulianov S V., et al. 2017. Single-nucleus Hi-C reveals unique chromatin reorganization at oocyte-to-zygote transition. Nature 544: 110–114.

Heinz S, Benner C, Spann N, Bertolino E, Lin YC, Laslo P, Cheng JX, Murre C, Singh H, Glass CK. 2010. Simple combinations of lineage-determining transcription factors prime cis-regulatory elements required for macrophage and B cell identities. Mol Cell 38: 576–89.

Imakaev M, Fudenberg G, McCord RP, Naumova N, Goloborodko A, Lajoie BR, Dekker J, Mirny LA. 2012. Iterative correction of Hi-C data reveals hallmarks of chromosome organization. Nat Methods 9: 999–1003.

Kent WJ, Sugnet CW, Furey TS, Roskin KM, Pringle TH, Zahler AM, Haussler a. D. 2002. The Human Genome Browser at UCSC. Genome Res 12: 996–1006.

Kim D, Pertea G, Trapnell C, Pimentel H, Kelley R, Salzberg SL. 2013. TopHat2: accurate alignment of transcriptomes in the presence of insertions, deletions and gene fusions. Genome Biol 14: R36.

Knight PA, Ruiz D. 2013. A fast algorithm for matrix balancing. IMA J Numer Anal 33: 1029–1047.

Krueger F, Andrews SR. 2016. SNPsplit: Allele-specific splitting of alignments between genomes with known SNP genotypes. F1000Research 5: 1479.

Langmead B, Salzberg SL. 2012. Fast gapped-read alignment with Bowtie 2. Nat Methods 9: 357–359.

Li H, Durbin R. 2009. Fast and accurate short read alignment with Burrows-Wheeler transform. Bioinformatics 25: 1754–1760.

Li H, Handsaker B, Wysoker A, Fennell T, Ruan J, Homer N, Marth G, Abecasis G, Durbin R, 1000 Genome Project Data Processing Subgroup. 2009. The Sequence Alignment/Map format and SAMtools. Bioinformatics 25: 2078–2079.

Pertea M, Pertea GM, Antonescu CM, Chang T-C, Mendell JT, Salzberg SL. 2015. StringTie enables improved reconstruction of a transcriptome from RNA-seq reads. Nat Biotechnol 33: 290–295.

Quinlan AR, Hall IM. 2010. BEDTools: a flexible suite of utilities for comparing genomic features. Bioinformatics 26: 841–2.

Ramírez F, Ryan DP, Grüning B, Bhardwaj V, Kilpert F, Richter AS, Heyne S, Dündar F, Manke T. 2016. deepTools2: a next generation web server for deep-sequencing data analysis. Nucleic Acids Res 44: W160–W165.

Robinson JT, Thorvaldsdóttir H, Winckler W, Guttman M, Lander ES, Getz G, Mesirov JP. 2011. Integrative genomics viewer. Nat Biotechnol 29: 24–26.

Servant N, Varoquaux N, Lajoie BR, Viara E, Chen C-J, Vert J-P, Heard E, Dekker J, Barillot E. 2015. HiC-Pro: an optimized and flexible pipeline for Hi-C data processing. Genome Biol 16: 259.

Zhang Y, Liu T, Meyer CA, Eeckhoute J, Johnson DS, Bernstein BE, Nussbaum C, Myers RM, Brown M, Li W, et al. 2008. Model-based Analysis of ChIP-Seq (MACS). Genome Biol 9: R137.

